# New analysis pipeline for high-throughput domain-peptide affinity experiments improves SH2 interaction data

**DOI:** 10.1101/2020.01.02.892901

**Authors:** Tom Ronan, Roman Garnett, Kristen Naegle

## Abstract

Protein domain interactions with short linear peptides, such as Src homology 2 (SH2) domain interactions with phosphotyrosine-containing peptide motifs (pTyr), are ubiquitous and important to many biochemical processes of the cell. The desire to map and quantify these interactions has resulted in the development of high-throughput (HTP) quantitative measurement techniques, such as microarray or fluorescence polarization assays. For example, in the last 15 years, experiments have progressed from measuring single interactions to covering 500,000 of the 5.5 million possible SH2-pTyr interactions in the human proteome. However, high variability in affinity measurements and disagreements about positive interactions between published datasets led us to re-evaluate the analysis methods and raw data of published SH2-pTyr HTP experiments. We identified several opportunities for improving the identification of positive and negative interactions, and the accuracy of affinity measurements. We implemented model fitting techniques that are more statistically appropriate for the non-linear SH2-pTyr interaction data. We developed a novel method to account for protein concentration errors due to impurities and degradation, as well as addressing protein inactivity and aggregation. Our revised analysis increases reported affinity accuracy, reduces the false negative rate, and results in an increase in useful data due to the addition of reliable true negative results. We demonstrate improvement in classification of binding vs non-binding when using machine learning techniques, suggesting improved coherence in the reanalyzed datasets. We present revised SH2-pTyr affinity results, and propose a new analysis pipeline for future HTP measurements of domain-peptide interactions.

## Introduction

Protein domain interactions with short linear peptides are found in many biochemical processes of the cell, and play a central role in cell physiology and communication. For example, SH2 domains are central to pTyr signaling networks, which control cell development, migration, and apoptosis (*1*). The 120 human SH2 domains are considered “readers”, since they read the presence of tyrosine phosphorylation by binding specifically to certain phosphorylated amino acid sequences. Approximately half of the binding energy of the SH2-pTyr sequence interaction is due to an invariant arginine which creates a salt bridge with the ligand pTyr. The remainder of the binding energy results from interactions between the SH2 domain binding pocket and the residues flanking central pTyr residues (*2*–*4*), resulting in specificity of SH2 domain interactions critical to pTyr-mediated signaling (*5*). Measurements of all SH2 binding affinities for target peptides would greatly aid in the decryption of domain specificity and advance understanding of cell signaling networks that control human physiology. However, the total potential number of interactions is immense – the 46,000 tyrosines currently known to be phosphorylated in the human proteome (*6*) have the potential to interact with 120 human SH2 domains, resulting in over 5.5 million possible SH2-pTyr interactions.

Recent developments have expanded the measurement coverage of human SH2-pTyr interactions. Eight high-throughput (HTP) studies have been performed to measure SH2 domain interactions with specific phosphopeptide sequences (*7*–*14*) (Table 1) using either microarrays or fluorescence polarization (FP). The six studies that quantitatively measured affinity represent ∼90,000 pairs of domain-peptide interactions, but these measurements cover less than 2% of possible interactions. In response, computational approaches have attempted to predict as-of-yet-unmeasured interactions using the published interaction data. These methods span the range from thermodynamic models which predict interaction strength using existing structure and binding measurements (*15*–*17*) to supervised machine learning models using patterns in peptide sequences and quantitative binding data to predict binding (*14*, *18*). However, no computational method has used the available affinity data in its entirety. We therefore wished to leverage all available binding affinity data in a supervised learning approach to expand our knowledge of SH2-pTyr interaction space.

**Table 1:** Overview of Published SH2 Data and Use in Published Models. Eight high-throughput experiments have been published since 2006 using experimental techniques such as protein microarrays (PM), peptide arrays (PA), and fluorescence polarization (FP). Of the published studies, only two studies have raw data available, by personal communication. Even the published data from several studies is no longer available. (pos) Raw data only published for positive interactions; (PC) data available only by personal communication; (fig) Published as a figure only, numerical results are available by private communication; (*) Original results were stored in PepspotDB, but not published in the journal or supplement. PepspotDB is no longer available.

| Data Group | Type | Publication | Peptides | SH2 Domains | Affinity | Results Available | Raw Data Available | Models |  |
| --- | --- | --- | --- | --- | --- | --- | --- | --- | --- |
| 1 | PM | Jones et al. (2006) | 61 | 159 | Yes | Yes | No | SH2PepInt | Wunderlich/Mirny |
|  |  | Kaushansky et al. (2008) | 50 | 133 | Yes | Yes | No |  |  |
|  |  | Gordus et al. (2009) | 46 | 96 | Yes | Yes | Yes (pos) |  |  |
| 2 | PM | Koytiger et al. (2013) | 729 | 70 | Yes | Yes | No | MSM/D, FoldX |  |
| 3 | FP | Hause et al. (2012) | 89 | 93 | Yes | Yes | Yes (PC) |  | PEBL |
|  |  | Leung et al. (2014) | 85 | 93 | Yes | Yes | Yes (PC) |  |  |
| n/a | PA | Liu et al. (2010) | 192 | 50 | No | No (fig) | No |  |  |
|  |  | Tinti et al. (2013) | 6202 | 70 | No | No (*) | No (*) |  |  |

Unfortunately, in the process of reviewing published HTP data, we found surprising disagreement between publications about which domain-peptide pairs interacted. For the limited number of interactions for which they agreed, they reported vastly different affinities for interacting pairs. We identified two issues common to all of the data sets that could be responsible for the discrepancies in results: errors affecting protein concentration, and improper use of statistical methods affecting modeling results.

First, we found potential sources of errors in protein concentration that could affect reported affinity values. Protein was minimally purified (via nickel chromatography), and protein concentration was measured by absorbance. No study used positive controls to determine the degree of protein functionality before measuring affinity. Thus protein of varying degrees of purity, functionality, and non-monomeric content were used for affinity measurements. Impure or degraded protein causes overestimation of protein concentration when compared to the amount of active protein in a sample. These protein concentration errors can propagate directly to errors in affinity, because affinity is derived from concentration and activity.

Second, we found errors in model fitting and statistical methods used to evaluate model fitting, which could have significant impact on the reported affinities. All of the affinity studies used the receptor occupancy model and the coefficient of determination (r^2^) as a determination of how well the model fits the data. For *linear* models, one can interpret the values of r^2^ between 0 and 1 as the total percent of variance explained by the fit. However, when applied to *non-linear* models (like the receptor occupancy model used in each of these studies to derive affinity), the r^2^ value cannot be interpreted as the percent of variance, and has been conclusively shown to be a poor indicator of fitness (*19*). Although this fact has long been established in the statistical literature (*20*–*26*) r^2^ is still commonly used to evaluate non-linear models in pharmaceutical and biomedical publications despite being an ineffective and misleading metric. In these publications, the use of r^2^ effectively resulted in a bias for true positive interactions at the expense of making many false negative calls, and removed many replicate measurements from incorporation into the reported affinity values.

Therefore, due to both inaccuracies in quantitative results and the significant potential for large numbers of false negative results, we had serious concerns about using the published affinities in machine learning. To overcome these issues, we decided to retrieve and reanalyze available raw data in order to systematically improve classification and affinity accuracy for SH2pTyr interactions. To accomplish this we: 1) refined model fitting techniques, 2) implemented fitting multiple models to each measurement, 3) used a statistically accurate method for model selection, 4) developed methods to identify and remove non-functional protein from the results, and 5) introduced a novel method to address the effects of protein concentration errors on reported affinity.

Our revised analysis improves affinity accuracy, improves specificity by reducing the false negative rate, and results in a dramatic increase in useful data due to the addition of thousands of true negative results. Evaluation of the revised affinities shows improved learning accuracy within an active learning model – suggesting that there is improved coherency in the features of the revised dataset.

## Results

### Evaluation of published affinity data and acquisition of raw data

In the process of evaluating published high-throughput data, we found significant disagreement between data sets. We evaluated all publications using HTP methods to measure SH2 domain interactions with specific peptide sequences. The publications containing SH2 affinity data can be grouped into three, distinct data groups (Table 1). The first data group consists of the group of studies published by the MacBeath lab from 2006 to 2009 (*7*, *9*, *27*) which contain a body of predominantly non-overlapping protein microarray (PM) experiments. The second data group consists of a large study published by the MacBeath lab in 2013 (*10*) with a set of new PM measurements using the protocol published in 2010 (*28*). The third data group consists of two non-overlapping sets of fluorescence polarization (FP) experiments published in 2012 and 2014 by the Jones lab (*13*, *14*). Because the other experiments (*11*, *12*) only measured interaction and not affinity, they were not considered for this analysis.

In order to determine agreement between data sets, we examined both qualitative and quantitative results. First, we examined the correlation between domain-peptide affinity measurements which overlapped between any two data groups (Fig. 1, top row). We found surprisingly low correlation between affinity measurements (with a maximum correlation of r = 0.367). Next we asked if the different data groups identified the same positive interactions between domain-peptide pairs, even if they did not agree on the affinity measurements. Here, we found significant disagreement over which domain-peptide pairs were found to interact (Fig. 1, bottom row). Of 347 positive domain-peptide interactions identified by one or more groups, fewer than 16% (55/347) were found to interact in all three data groups. No two experiments were able to agree on more than 29% of the positive interactions. The differences in interaction identification were spread randomly among SH2 domains and peptides, with no single SH2 domain, peptide, or peptide family being overrepresented (Fig. S1). These findings demonstrate significant quantitative and qualitative differences between published data from different labs, and even disagreement between publications from the same lab.

**Figure 1:**
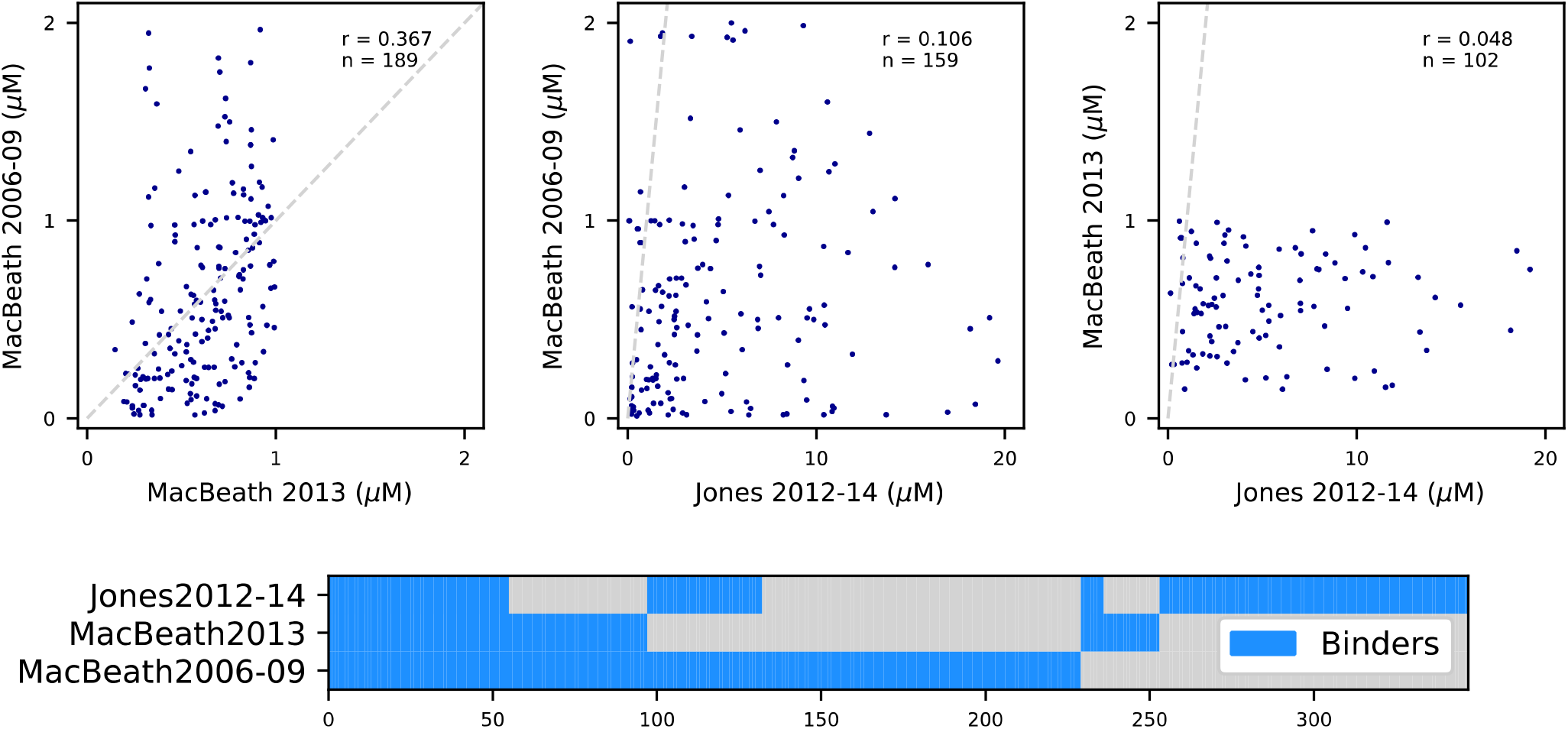
Comparison of Published Affinity Data. Correlation of affinity from three data groups is evaluated using scatter plots (top row). See Table 1 for group definitions. With perfect agreement, data points would fall along the dashed gray line. Surprisingly, there is very low correlation between affinities from different data groups. Even results from the same lab published at different times show only mild correlation (r=0.367, MacBeath 2006-09 vs MacBeath 2013). The data were also examined for agreement on positively interacting domain-peptide pairs (bottom panel). Positive interactions are identified by blue bars. Of the 347 positive domain-peptide interactions identified by at least one group, only 55 interactions were found to be positive in all three data groups (15.9%). No two data groups agreed on more than 29% of positive interactions. Although there are significant differences between the techniques of protein microarrays (PM) and fluorescence polarization (FP), the differences between identities of positive interactors did not segregate by experimental technique. The two PM experiments (MacBeath 2013, MacBeath 2006-09) identified 28.0% (97/347) of the positive interactions in common, but a similar number of positive interactions can be seen (25.9%, 90/347) when comparing one PM experiment with the FP experiment (MacBeath 2013, Jones 2012-14). The highest correlation was also between experiments of different techniques ((r=0.367, MacBeath 2006-09 vs MacBeath 2013).

Although there are significant differences between the techniques of PM and FP experimental methods, the differences between positive interactors did not group by technique (Fig. 1) and different techniques had similar numbers of common positive interactions. Although very low, the best correlation (r = 0.367) was between a PM experiment and an FP experiment from the same lab. All three data groups used similar protein and peptide production and purification methods, absorbance for determination of protein concentration, the receptor occupancy model for determining affinity, and similar methods of evaluating model fits based on the coefficient of determination (r^2^).

We concluded that we would need to look further into the methods and raw data to evaluate the differences between published data sets, or even to evaluate the quality of any single set of published data. Acquisition of raw data from published studies was surprisingly difficult. No publication included raw data, only supplemental tables with post-processed values for affinity which are insufficient for replication of published results. Furthermore, we discovered that most raw data underlying the published analysis has been lost by the original authors and is no longer available from any party. (Table 1) Fortunately, we were able to retrieve raw data from the Jones 2012-14 data group (personal communication from Richard Jones, Ron Hause, and Ken Leung).

### Raw SH2 interaction data and revised analysis

We began by examining the raw data from the Jones 2012-14 data group to evaluate the quality and completeness of the data, and to review the methods used to process the raw data into its published form. Although some raw data was missing in comparison to the original publication, by limiting our revised analysis to interactions of single SH2 domains with phosphopeptides from the ErbB family (EGFR, ERBB2, ERBB3, ERBB4), as well as KIT, MET, and GAB1, the available raw data covered approximately 99.6% of the reported measurements.

#### Evaluation of the implementation of the receptor occupancy model

The raw data for each measured interaction consisted of fluorescence polarization measurements of an SH2 domain in solution with a phosphopeptide at equilibrium at 12 concentrations. In the original publication, the raw data was then used to derive an equilibrium dissociation rate constant (K_d_) by fitting the receptor occupancy model (developed by Clark in 1926 using the law of mass action (*29*)). As applied to the fluorescence polarization data, the model takes the form:

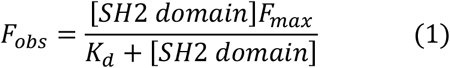

where F_obs_ is the observed FP signal at each assayed protein concentration of the SH2 domain (measured in millipolarization units (mP)), and F_max_ represents the FP at saturation (see also Fig. S2). The affinity (K_d_) and saturation limit (F_max_) are fitted parameters of the model. It is important to note that this model is dependent on several critical assumptions: that the reaction is reversible; that the ligand only exists in a bound and unbound form; that all receptor molecules are equivalent; that the biological response is proportional to occupied receptors; and that the system is at equilibrium.

We hypothesized that the specific methods used to implement the receptor occupancy model in the original publications might have affected the accuracy of the originally published results. We examined three aspects of the implementation of this model. First, we reviewed the method of subtraction of background fluorescence and found that it introduces systematic random errors in affinity results. Second, we evaluated whether the receptor occupancy model could reliably fit a non-binding sample. When we found it could not, we implemented an alternate model and a model selection procedure in order to more reliably identify negative interactions. Finally, we examined the effect of dropping outlier measurements on model fitting results, and implemented an alternative method to determine model fitness: signal-to-noise ratio (SNR).

#### Background subtraction causes errors in model fits and is replaced by fitting an offset

In the original analysis, the authors used a plate-wise background subtraction method, where the median baseline control value was recorded from plate measurements and subtracted from the polarization signal observed at each data point (*13*). When plates had excessive variation in baseline control values, the authors excluded these results from further analysis. However, in examining many measurements by eye, we found that the background values seemed uncorrelated with the signal values (Fig. S3).

A critical feature of the receptor occupancy model is that the saturation curve passes through the origin (because the point of zero-signal is also the point of zero-concentration). Thus background subtraction forces the zero-signal to a point other than that of zero-concentration, resulting in higher residual error which introduces errors in derived affinity (Fig. S4). These errors increase or decrease affinity (based on whether the background is high or low), and are non-linear by affinity. Since the relationship of the background was seemingly random, and the error factors are non-linear, background subtraction injected random error in affinity calculations. More than 54% of the replicate measurements exhibited problematic background levels (Fig. S4, bottom row). Thus, we rejected the background subtraction method in favor of fitting the model along with an offset value (F_0_):

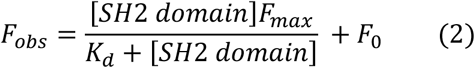

Example fits using the receptor occupancy model with offset can be seen in Fig. S5.

#### The receptor occupancy model fails to accurately identify non-binding measurements; introducing a linear model

Although the receptor occupancy model is theoretically capable of fitting a typical binding saturation curve as well as a ‘flat’ curve representative of non-binding interactions, we found that it fails to accurately fit non-binding interactions in practice (Fig. S6), resulting in artefactual model fits with unreasonable affinity and saturation values.

Because negative interactions resemble low-slope or zero-slope lines with superimposed random noise, we hypothesized that a linear model would more reliably fit these ‘non-binders’ and resolve fit artifacts. The linear model is:

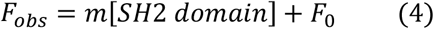

(where F_0_ represents an offset value, and *m* is a constant representing the slope of the fitted line, Fig. S6, red fits). Out of 37,378 replicate measurements, 31,861 were best fit by the linear model. Of these, 29,778 were initially classified as non-binders.

We also found a group of replicate measurements (∼6%) which were best fit by a linear model but with steep positive slope. Linearly increasing fluorescent signal with no indication of saturation violates the assumptions of a receptor occupancy model, and is more likely to represent a form of protein aggregation, peptide aggregation, or some other form of non-specific binding. Thus, to preserve the quality of the non-binding calls, a conservative slope cutoff of 5mP/μM was implemented, above which replicates were identified as aggregators, and removed from further consideration. Of the 31,861 replicates best fit by the linear model, 2083 were initially classified as aggregators.

#### Fitting multiple models requires a model selection process

When more than one model can be used to fit the data, a method of model selection must be implemented to determine which model most accurately represents the data while balancing against adding additional parameters which can lead to overfitting. In order to determine if a measurement is best described by a receptor occupancy model or a linear model we used the Akaike Information Criterion (AIC). In contrast to the coefficient of determination (r^2^), AIC is a model selection metric which is appropriate for use with non-linear models (*19*, *30*), is robust even with high noise data, and employs a regularization technique to avoid overfitting by penalizing models with more parameters (the receptor occupancy has 3 parameters; the linear model has only 2 parameters). In our implementation we used a bias corrected form of the metric, AICc, in order to account for only having 12 data points per saturation curve. A lower AICc score indicates a better fit. Examples of model fitting can be seen in Fig. S7.

#### Evaluation of model fitness

In order to determine how well the data was represented by a model, we used the signal to noise ratio (SNR) as a metric of model fitness. The SNR metric represents the magnitude of residual errors of fit to the model (a form of noise), and weighs this sum by the overall size of the fluorescent signal measured. It is calculated as

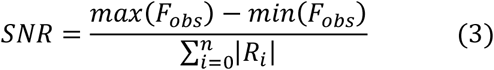

where *n* is the number of data points, R_i_ is the residual value of the *i*^*th*^ data point, and F_obs_ is the observed fluorescence (in mP units).

At an SNR≥1, the measured signal is larger than the sum of all errors to the fit, and represents a good quality fit in practice. We chose a ratio of 1 as the limit of a good fit based on extensive visual inspection of the fits (see Fig. S8 and Fig. S9). Replicates with SNR<1 made up 5.2% of fits (1948/37378). These low-SNR fits fell into three classes: zero signal measurements (76.0%, 1480/1948), measurements where noise swamped the signal (21.9%, 427/1948), and good measurements with large single point outliers (2.1%, 41/1948). Although the metric excludes some viable measurements that would be kept when reviewing by eye (e.g. 41 single point outlier replicate measurements) this represents only 1/10th of 1% of all replicate measurements (0.11%, 14/37378). A consistent standard is difficult to implement without such a metric, and any objective metric would likely excluse some viable measurements. (See the Discussion for thoughts on alternate metrics.)

#### Outlier removal biases model fitting and selection

In the original publications, the authors utilized an iterative outlier removal process. For each set of 12 data points in a replicate measurement, individual points were identified as outliers using a statistical model. The outlier was removed and the fit was reevaluated. Up to three points were removed per replicate measurement. For measurements where more than three data points were identified as outliers, the replicate measurement was removed from further consideration.

For an ideal binding saturation experiment attempting to identify K_d_, the concentrations tested should span either side of K_d_, and the highest and lowest measured concentrations should establish the plateaus seen on semi-log saturation plots (See Fig. S5). Based on the concentrations selected for this experiment, the ideal range for quantification is affinity (K_d_) in the range of 0.05μM to 0.5μM. For interactions with a K_d_>1.0μM, the upper plateau of the semi-log saturation curve no longer has any coverage (Fig. S5, row 2). Interactions with K_d_>5μM have no data points at all above K_d_ (Fig. S5, rows 3 and 4). This suggests that every data point is critical for accuracy, particularly points above K_d_, thus we chose to use all data points to avoid introducing additional error and to allow the SNR metric to gauge the quality of fit.

#### Summary of revised analysis method for replicate measurements

Following a systematic review of each decision made in evaluating a measurement in HTP affinity studies we developed an improved analysis pipeline (Fig. 2). For each replicate measurement, we fit two models: a receptor occupancy model with offset (equation 2) and a linear model with offset (equation 4). Fits were evaluated with AICc: the model with the lower score was chosen as the best fit. Replicates that were fit best by the linear model and had a slope of less than or equal to 5mP/μM were classified as negative interactions, or ‘non-binders’. Linear fits with a slope greater than 5mP/μM were classified as aggregators and removed from considerations. A replicate that was fit best by the receptor occupancy model was then evaluated for signal-to-noise ratio (SNR). If the SNR was greater than one, the replicate was classified as a positive interaction or ‘binder’. Out of 37,378 replicate measurements, we identified 7.4% (2753) as binders, 79.7% (29,778) as non-binders, 7.4% (2764) as low-SNR fits, and 5.6% (2083) as aggregators (Fig. 3, left side).

**Figure 2:**
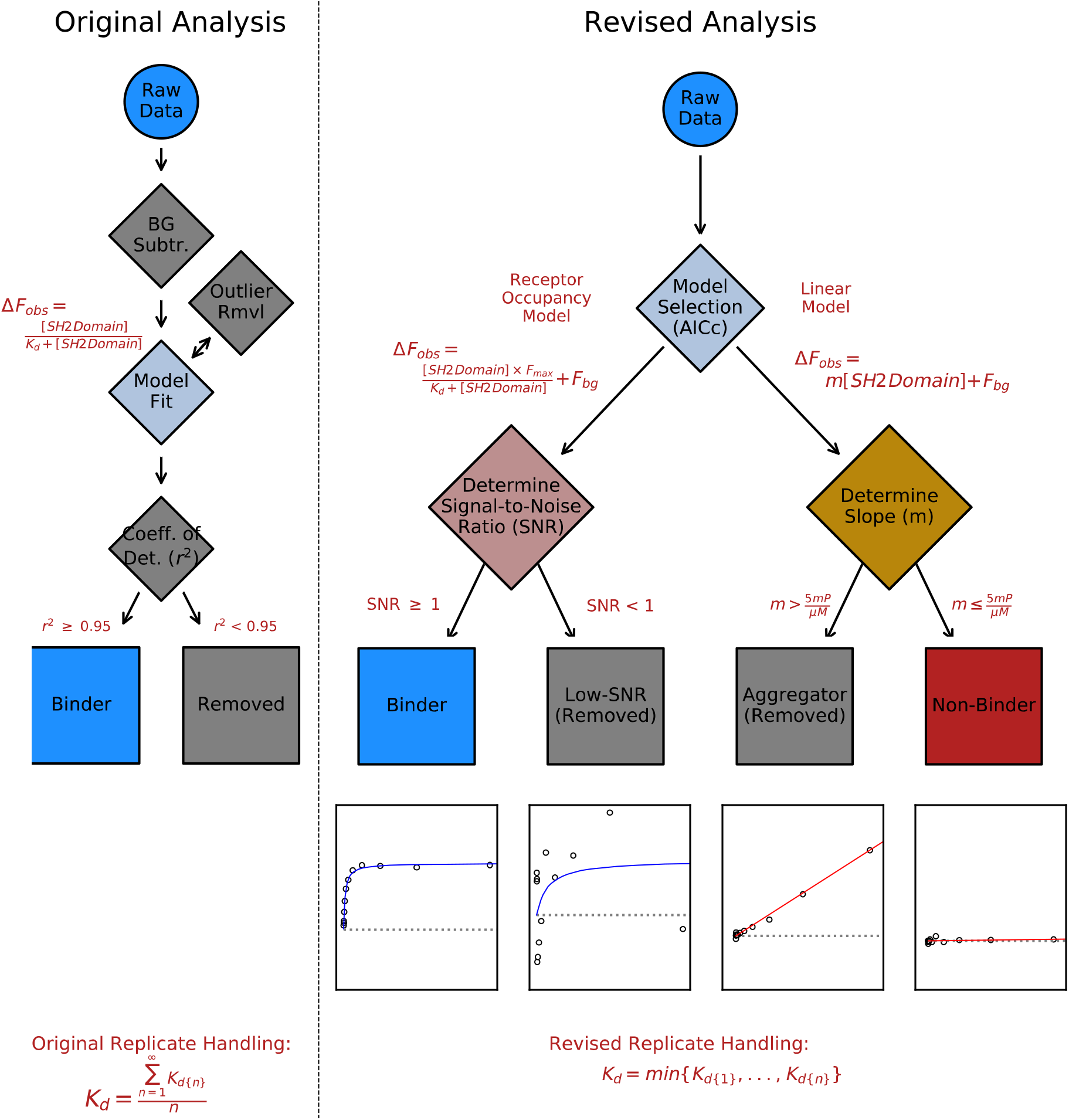
Flowchart of Revised Analysis Process. Comparison between the original analysis process (left panel) and our revised analysis pipeline (right panel). For our revised process, representative sample fits are shown below each of the final categorizations.

**Figure 3:**
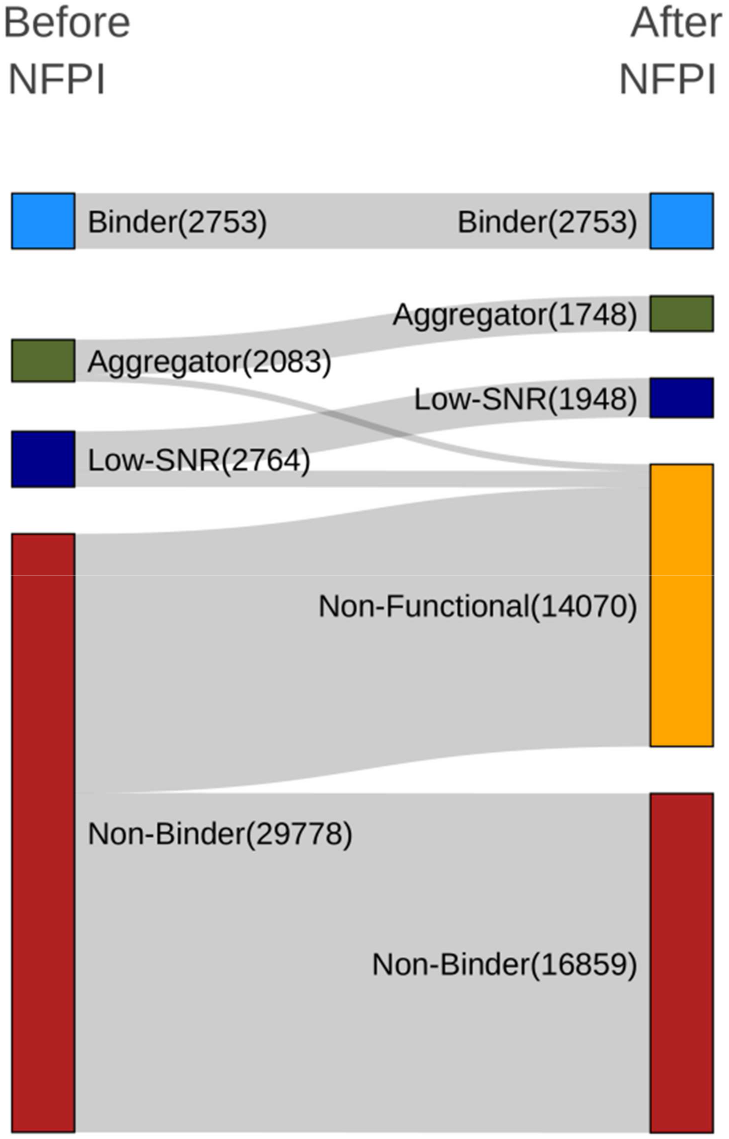
Initial Replicate-Level Results and the Results of Non-Functional Protein Identification (NFPI). The categorization results of individual domain-peptide measurements are shown (Before NFPI). Of the 37,378 measurements, 7.4% (2,753) were initially identified as positive interactions (binders), 7.4% (2,083) as interactions showing aggregation, 5.6% (2,764) as low signal-to-noise, and 79.7% (29,778) as non-binders. The subsequent identification and removal of individual domain-peptide measurements made on non-functional protein had a significant effect on the categorization of non-positive replicate-level measurements. Of the 29778 measurements initially categorized as non-binders, 56.6% (16,859) were identified as likely to contain non-functional protein and were removed from further consideration.

### High variation at the replicate level is likely caused by protein concentration errors

The original publication reported a single affinity (K_d_) value for each domain-peptide pair, which was the average of multiple replicate domain-peptide measurements. However, we found patterns of high variation in affinity between replicates that suggested a significant problem with the either experimental design or experimental method. We hypothesized that a single variable – errors in protein concentration – could be responsible for the high variance.

In reviewing data quality, we identified a large number of domain-peptide interactions demonstrating high variance in affinity among replicates (for example, K_d_ values ranging from below 0.5μM to over 20μM for replicates from a single domain-peptide interaction). In order to determine the source and character of the variance, we inspected replicates as a group for individual domain-peptide interactions (for a representative example, see Fig. S10). Despite high variance between replicates, each replicate measurement had high quality fits and low residual error, as expected from meeting an SNR>1. (For a representative example of all measurements from one such replicate group, see Fig. S10).

To explore this further, we visualized and quantified variance (Fig. 4) for all domain-peptide interactions. Although variance tends to increase as K_d_ increases, variance greater than 10μM is found across a large fraction of all measurements, independent of affinity. How could high-quality individual replicate measurements result in such varied affinities for a single domain-peptide pair? We hypothesized that protein concentration error (arising from differences in protein preparations such as impurities, degradation and inactivity) could directly propagate to errors in modeled affinity values while still producing high-quality individual replicate saturation curves.

**Figure 4:**
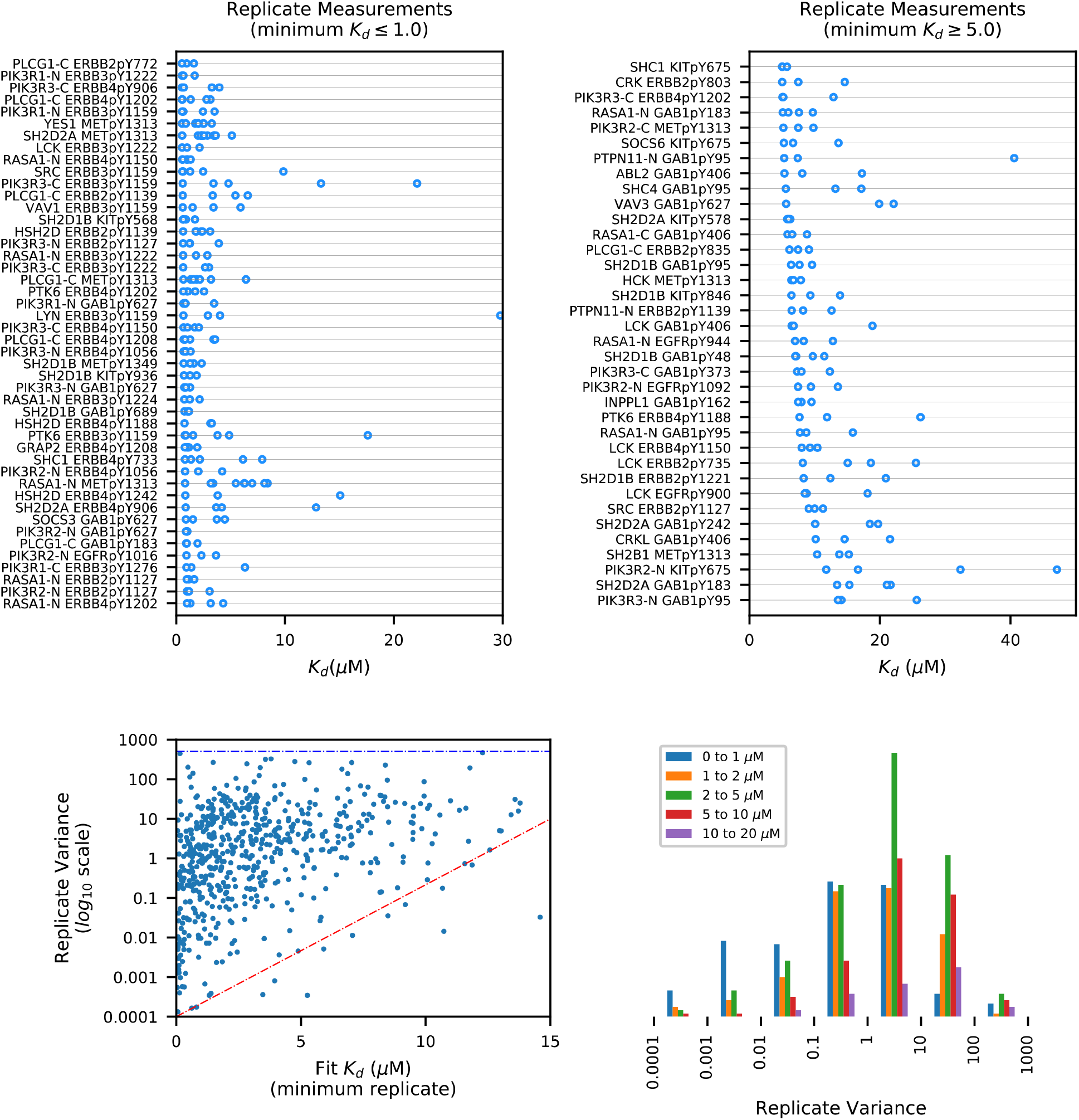
Replicate Measurements Exhibit High Variance. Variance in affinity for domain-peptide interactions was visualized by distributed dot plots (top row) for examples of higher-affinity (K_d_ ≤ 1.0μM, upper left) and lower-affinity (K_d_ ≥ 5.0μM, upper right) interactions. Each row displays all replicate measurements for a single domain-peptide pair, and the x-axis position of each individual replicate reflects the K_d_ value of that measurement. Thus variance can be visualized as the width of spread of points along each line. Domain-peptide interactions are sorted by minimum replicate K_d_. The relationship between variance and affinity was also visualzed for all domain-peptide interactions (lower left, note the y-axis log-scale). The minimum replicate variance generally increases as K_d_ inceases (trend indicated by the red-dashed line) but worst-case variance is independent of K_d_ (blue-dashed line), and high variance (e.g. ≥ 10) is present at all K_d_ ranges. Variance was also quantified for different minimum K_d_ ranges against different variance ranges (lower right). In extremely low variance cases (e.g. variance ≤ 0.01), low K_d_ measurements (blue bar) dominate. In moderate to high variance ranges (e.g. ≥ 1), the distributions are more similar. These two trends support the reasonable inference that higher K_d_ fits have higher variance in general, but also demonstrates that the presence of high variance replicates is independent of affinity.

To test this hypothesis, we first examined the theoretical effects of protein concentration error on affinity. We demonstrate that concentration errors directly manifest as errors in affinity, and that errors from impurity or degradation systematically manifest as artificially high K_d_ (lower affinity). Next, we examined the methods and data for sources of purification errors, partial degradation, and complete protein inactivity, and identified evidence of all three. Finally, we developed a method to control for these sources of protein concentration error and produce affinities with higher accuracy using the existing raw data.

#### Protein concentration errors propagate directly as errors in derived K_d_ values

Although binding affinity is a molecular property – affinity is the strength of interaction between a single protein molecule and a single peptide – accurate derivation and calculation of affinity by most methods depends on the accuracy of concentration measurements for the tested protein. In the case of the receptor occupancy model used here, affinity is a derived function of concentration and FP response. Because impurities or degraded protein represent an error between the assumed concentration and the active concentration of a protein, we hypothesized this would propagate to errors in affinity.

We examined the theoretical effect of concentration errors on measured affinity (Fig. 5). Errors in protein concentration due to impurities or degradation cause an overestimation of the true concentration of active protein. Overestimation errors in protein concentration cause errors in K_d_, always resulting in a higher K_d_ (lower affinity) than the true value. This error is linearly proportional to the error in concentration.

**Figure 5:**
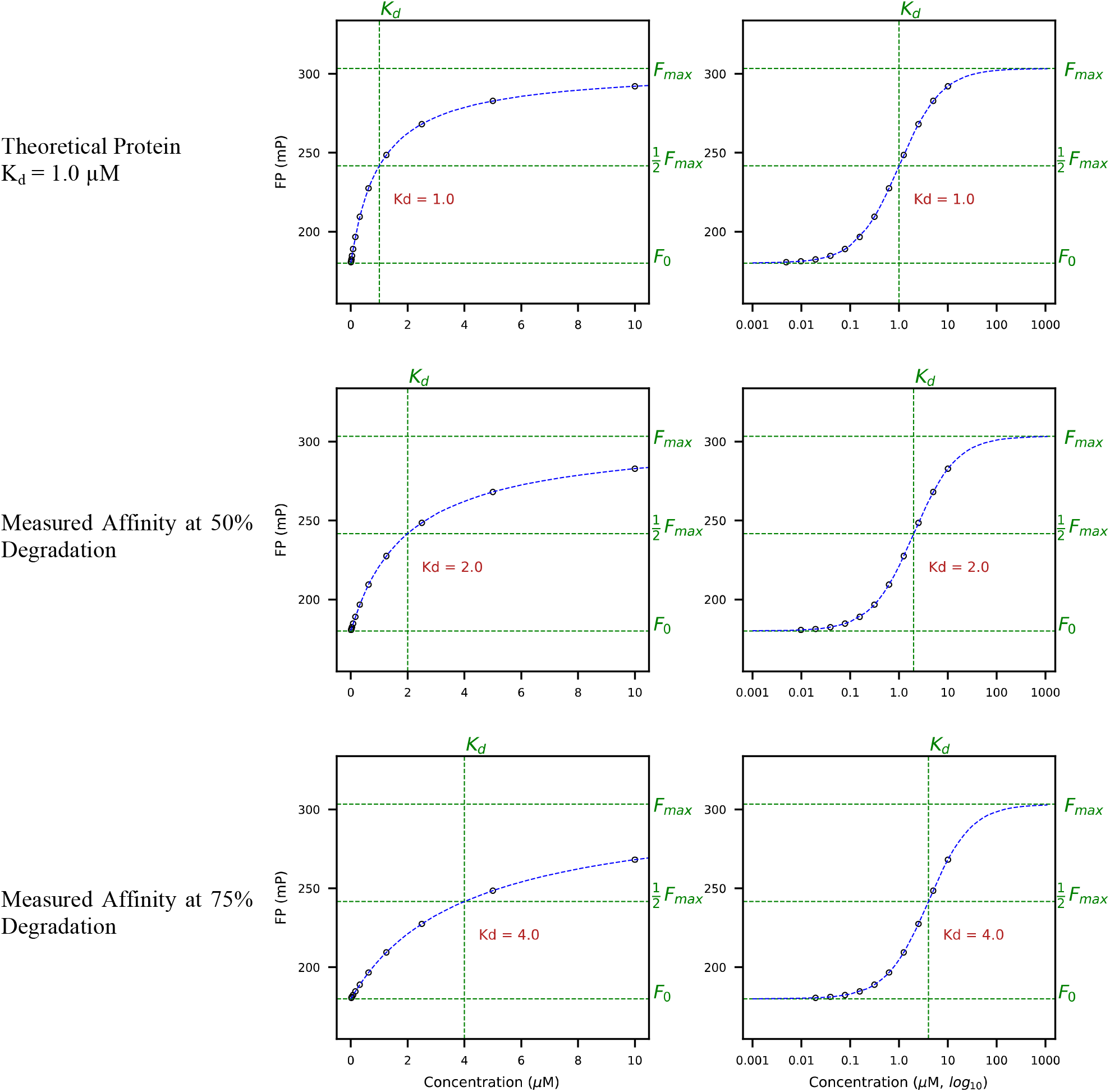
Degradation Causes a Decrease in Measured Affinity. Simulated measurements for an ideal binding saturation experiment are shown for a theoretical protein with a K_d_ of 1 μM (row 1). (Measurements in the second column are the same data as the first column, but plotted as semi-log plots with a logarithmic concentration axis.) In rows 2 and 3, the results of 50% and 75% degradation are shown. To simulate the effect of degraded protein, we plotted true activity from the ideal curve (row 1) against the erroneous assumed concentration due to degradation (rows 2 and 3). For example, to simulate 50% degradation (row 2), the true FP response for 5μM (from row 1) is plotted at the 10μM position (on row 2). This procedure is repeated at each concentration. When affinity is derived from these degraded protein measurements, the result is an inaccurate K_d_ higher than the true value. Although the concentration error from degraded protein causes a non-linear change in FP, the error in K_d_ is linear and proportional to the concentration error. For example, if the true active protein concentration is ½ of the assumed concentration (as in row 2), the measured affinity is ½ of the correct value (meaning the K_d_ is 2x the true value). If the true active protein concentration is ¼ of the assumed concentration (as in row 3), the measured affinity is ¼ of the correct value (the K_d_ is 4x the true value). Therefore errors from overestimation of protein concentration always result in higher measured K_d_ than the true value.

Thus, protein concentration errors due to batch impurities or degradation can manifest as a range of K_d_ values in replicate measurements made from different batches of protein, all of which would be equal to or higher than the true K_d_, while simultaneously coming from high-quality, low-noise replicate fits. This exact phenomenon has also been demonstrated experimentally (*31*).

#### Evidence for protein concentration errors due to protein degradation or impurity

The original publications used His_6_-tagged recombinant SH2 domain protein production methods, and used nickel chromatography as the sole protein purification method. In theory these methods can provide purities of up to 95% (*32*). However in practice the results can vary significantly, and can be affected by the amino acid content, nonspecific binding, purification conditions, and the type of affinity matrix used (*32*). Our experience in the lab performing these purifications suggests that differences in purity between different protein preparations are likely to be present. Because the method used to determine protein concentration was absorbance at 280nM, only total protein content is measured, independent of purity or activity.

If the variance in affinity was from a random (non-systemic) source, we would expect to find no patterns of variance in time. In contrast, if variance was from batch-related protein degradation or impurities, we might see alternating patterns in affinity over time as different batches are used. For example, if high-purity protein sample were used on run 1 and a low-purity protein sample were used on run 2, we would expect consistently higher affinities on run 1 and consistently lower affinities on run 2. Or, if a partially-degraded protein sample was exhausted mid-run, and replaced with a fresh sample, we could see a sudden surge of higher affinity results in the middle of a run, when compared to other runs. Similar patterns could arise from batch to batch variations in purity affecting accuracy of expected concentration.

We examined the data for evidence of these patterns. Since we do not have true information at the batch level or activity of each protein sample, these patterns must be inferred from the data. Although these patterns are difficult to spot due to the nature of the experimental design, we find examples of non-random run-dependent variations in affinity in the data (Fig. S11). These patterns are not compatible with a random source of variance, and are compatible with either degradation or protein impurity causing errors in protein concentration.

#### Evidence for complete non-functionality of protein domains

Because we found patterns consistent with partial degradation, we examined the data for patterns of complete protein degradation. Complete degradation, or completely non-functional protein, would be indistinguishable from a non-binding measurement for a single replicate, potentially resulting in a false-negative. A control experiment to determine protein functionality would normally be required to delineate these two cases. However, we hypothesized that non-functional protein would manifest within the data as long runs of non-binding results across many replicates, but would demonstrate contradictory evidence of binding on other runs when the protein was not degraded. We found patterns consistent with non-functional protein (Fig. S12). Non-functional protein domains were identified and removed from consideration (See Methods, and Fig. S13 and Fig. S14).

By removing replicates where there is evidence that the protein was non-functional, we avoid the potential for false negatives from this ambiguous data, and greatly improve the pool of true negative calls. Removal of non-functional protein has a significant impact on the numbers of measurements at the replicate level. Non-functional replicates made up 37.6% of all replicates (Fig. 3, right side). The large number of runs showing patterns of completely non-functional protein contributes to the overall evidence that protein degradation is present and is a source of variance in the data.

### Method for handling replicates with high variance due to protein concentration errors; Reporting the minimum instead of the mean

Two key issues arise when considering how to handle replicate measurements when impurities or degradation are suspected to be a primary source of variance. First, without knowing the exact amount of protein concentration error in any one sample, how can this error be controlled for? Second, what is the correct procedure for handling replicates when variation is primarily due to concentration errors and not random sample variation? We propose a simple but novel, solution to both questions: reporting the minimum rather than the mean of the replicate measurements results in the most accurate reported measurement.

Impurities and degradation can be partially controlled for by reporting the minimum replicate K_d_. Given some unknown amount of protein concentration error due to degradation or impurities, the *active* concentration of protein will always be equal to or lower than the *measured* concentration. And as we demonstrated above, this means that the true affinity of the protein will always be equal to or greater than the measured affinity. Put in terms of K_d_: the true K_d_ will always be equal to, or lower than the minimum measured K_d_. Thus, the minimum K_d_ reflects the closest measured value to the true affinity.

Furthermore, reporting the minimum measured K_d_ also addresses the variance problem. If the measurements were true replicates, reflecting random noise and experimental error, taking the mean of multiple replicates would be the appropriate procedure because the sample mean would represent the highest likelihood of the true population value of affinity. However, if the variation is caused by protein concentration errors, taking the mean of multiple measurements would not reflect the true affinity. Rather, it would inadvertently increase the reported K_d_ value by some unpredictable amount, which depends on the number of samples and the magnitude of their degradation. In addition, since the mean is particularly affected by outliers, even one severely degraded sample would significantly increase the mean reported K_d_ value, resulting in a reported affinity with high error. Therefore, odd though it may seem from a statistical perspective, taking the *minimum* K_d_ is the most accurate way to handle variation in replicates where errors in protein concentration overestimation represent the primary source of variation.

### Revised affinity results and comparison to the original published results

In the results from our revised analysis, 1518 positive (binding) interactions were identified, along with 7038 negative (non-binding) interactions. These ∼7000 true negative results represent a significant increase in information from the original publication in which no true negative interactions were reported. For 3200 interactions, inconclusive or problematic data was present and no conclusions about their affinity could be drawn. Of those, 2753 potential domain-peptide interactions remain unevaluated due to non-functional protein. Final affinity values were plotted for all peptide-domain interactions as a heat map (Fig. 6), and summarized by category of interaction and changes in calls (Fig. 7). A summary of our revised results and the originally published results are available in Supporting Data, as an Excel file, and the complete raw and revised data is available on Figshare (DOI: https://doi.org/10.6084/m9.figshare.11482686.v1).

**Figure 6:**
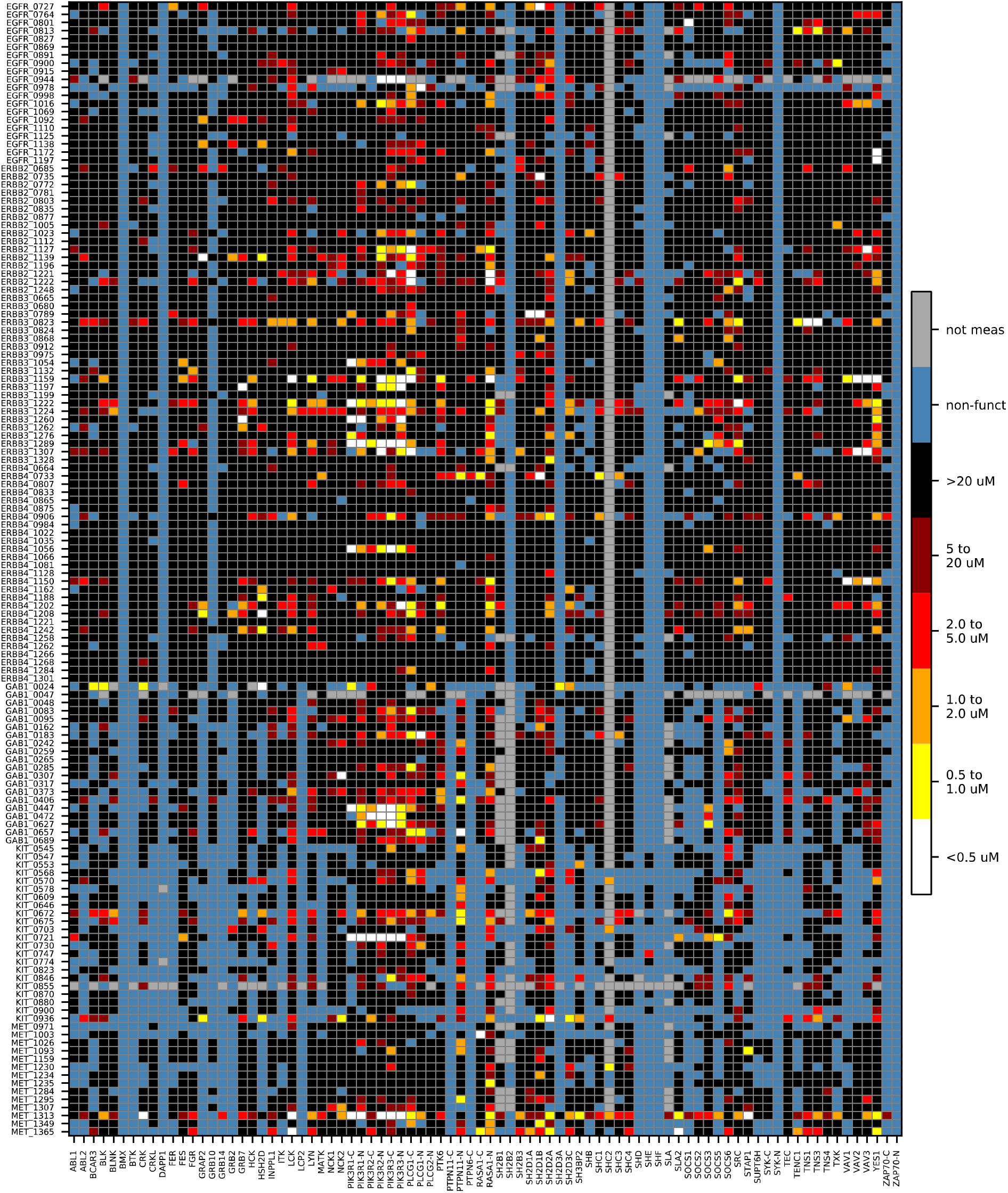
Revised Analysis Final Results. A heat map showing the final results of the revised analysis. A significant fraction a measurements demonstrated patterns consistent with non-functional protein and were removed from the analysis. Comparison with the original published results can be seen in Supplemental Figures 14 and 15.

**Figure 7:**
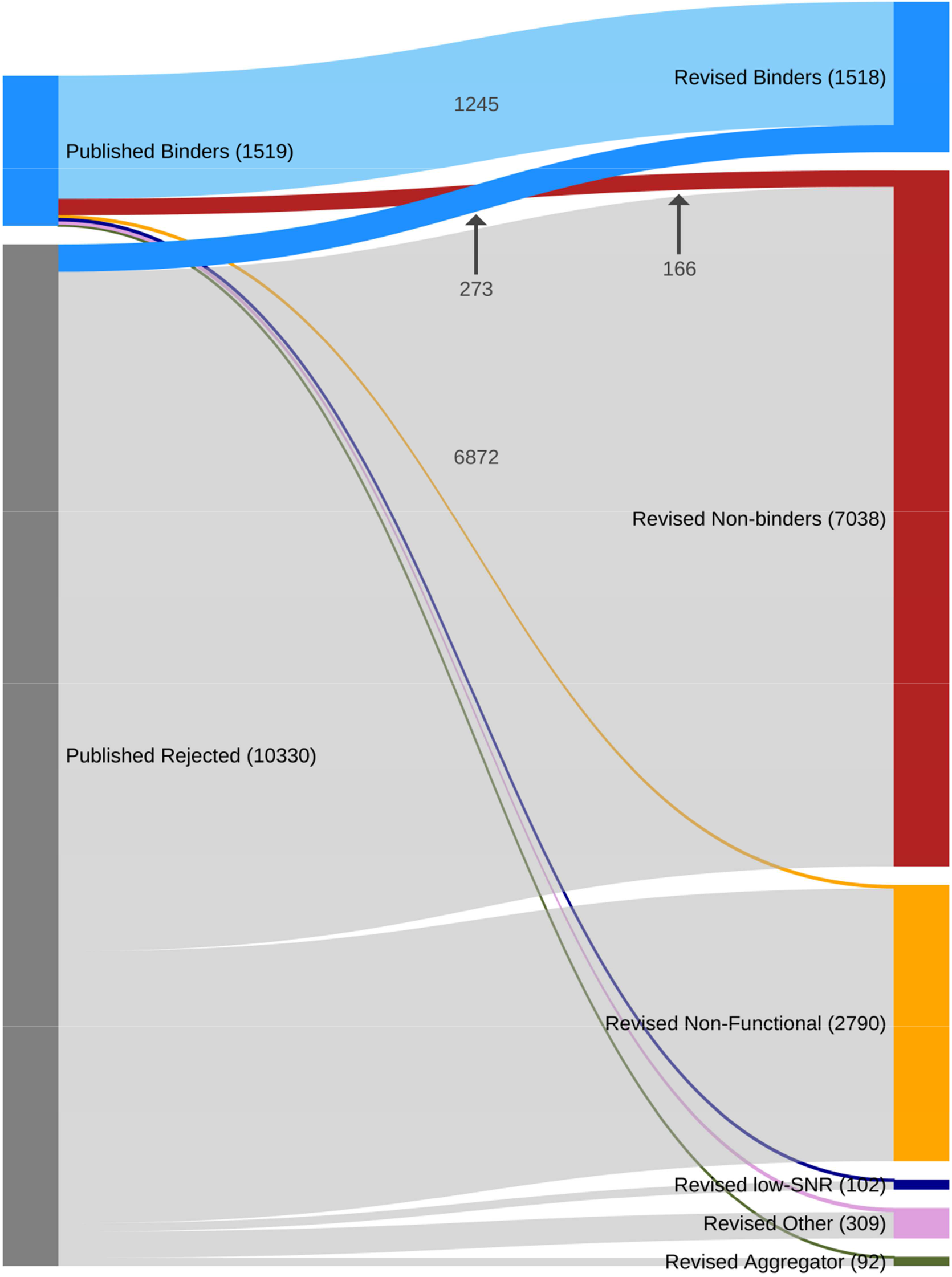
Changes In Calls Between Original Publication and Revised Analysis. Although the numbers of positive interactions are similar in our revised analysis, the identities of those interactions have changed significantly. The changes in calls are visualized in the Sankey map above. Of the original 1519 positive interactions found by the original authors, 166 (10.9%) were found to be non-binders in our analysis. Of the 10330 rejected interactions from the original publications, 273 (2.6%) positive interactions were recovered in our analysis.

Despite similar numbers of positive interactions between the original and revised results (1519 vs. 1518), the identities of the domain-peptide pairs comprising the positive interactions changed significantly (Fig. 7). More than 17% of the original positive interaction calls changed to either non-interactions, or rejected results due to data quality issues. In the final model, 166 interactions originally called positive in the published results are found to be true negative interactions. These changes are primarily due to the ability to avoid fit artifacts and false-positive results, a consequence of using multiple models to fit the data. Similarly large changes were found in the originally published negative interactions where 273 formerly rejected interactions are now classified as true positive interactions. These recovered results are primarily due to changes using offset fits instead of background subtraction, and using an appropriate quality metric to determine which model fits best. Changes in calls by class are visualized in Fig. 7, while the identities and magnitude of the domain-peptide pairs with changed calls are visualized in Fig. S15. Results from the original publication are visualized in Fig. S16.

Furthermore, even though 1245 domain-peptide pairs were found to bind in both the original publication and our revised analysis, the quantitative affinity of those binders changed significantly in the revised analysis (Fig. 8). Note that although the minimum of each replicate group was selected as most accurately reflecting the true affinity, our revised affinity values are not all lower than the original publication. This is primarily due to significant changes at the replicate level – where some original replicates were removed from consideration by changes in the fitting process, and a number of new replicates were included in each replicate set.

**Figure 8:**
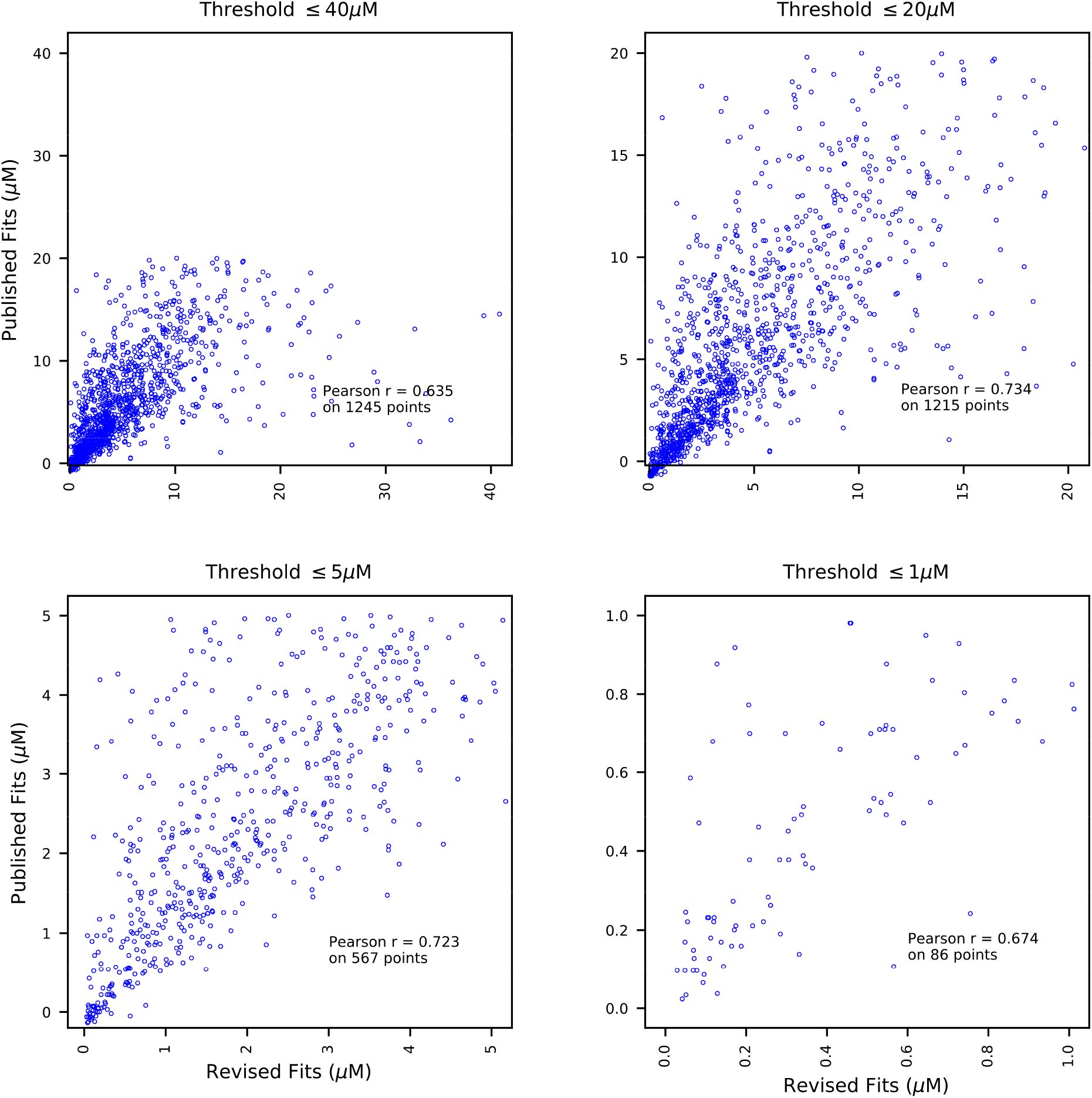
Correlation between Original Publication and Revised Analysis. Affinity values were compared for for the common set of positive interactions (n=1245, upper left panel), as well as at lower affinity thresholds (other panels, as indicated). Our revised affinity values correlate only moderately with the original publication (Pearson r=0.635), which might be surprising considering the analysis is on the same raw data. Our revised results correlate best when considering all measurements under 20μM affinity (Pearson r = 0.734). Despite choosing the minimum measured value for K_d_, our revised data often reports higher K_d_ results than the original publication (i.e. results below the diagonal). This is due to different categorization and filtering procedures which result in significant additions and removal of individual measurements in each set of replicates for a domain-peptide pair. It is interesting to note that correlation does not improve at higher affinity (lower K_d_), despite the fact that the chosen raw measurement range is tailored for highest accuracy for K_d_ < 1.0 μM. This suggests that the differences between our revised results are independent of the accuracy of the original measurements, and more likely due to the need to correctly handle variation due to protein concentration errors.

### Independent evaluation of revised analysis: measuring improved consistency via active learning

We wanted to evaluate our revised analysis compared to original results. In a case such as this, it is difficult to evaluate because original samples are no longer available. However, one way to evaluate the data is to use machine learning methods to ascertain whether the revised data has better internal consistency or predictive power than the original data set. Lacking a biological reference, it seemed fitting to evaluate this data using machine learning, as we originally wished to harness SH2 domain binding measurements in machine learning frameworks to extrapolate from the relatively small number of available measurements.

To do this, we implemented active search, a machine learning approach that is highly amenable to biochemistry problems such as this. Active learning (also known as optimal experimental design or active data acquisition) is a machine learning paradigm where we use available data to select the next best experiments to maximize a specific objective. Active search is a realization of this framework where the objective is to recover as many members of a rare, valuable class as possible. In this case where only 13.9% of the original dataset represents positive interactions between an SH2 domain and a phosphopeptide (or 18.2% in the revised dataset) the objective of the search algorithm was to prioritize each sequential selected interaction to maximize the total number of positive interactions discovered. We implemented the effective nonmyopic search (ENS) algorithm (*33*) with the goal of optimizing the total positive experiments identified in an allocated search of 100 queries. The algorithm was seeded randomly with one example positive before search progressed and was repeated 50 times.

ENS showed improved average performance and higher consistency with our revised dataset. First, ENS worked effectively on both the original and revised datasets, identifying positives that far exceed the expected number by random chance by the 100th query (Fig. 9). This suggests that phosphopeptide sequences do encode information about whether an SH2 domain will recognize them in a binding interaction. Second, ENS performance in the revised dataset was higher than the original dataset on average, finding 45.3 positives vs. 33.3 positives (p-value of 4e-12). Third, ENS performance is significantly more variable on the original dataset than on the final dataset (ranging between 9 and 62 positives in 50 trials (with an average of 33.3), compared to a range of 38 to 67 (with an average of 45.3 positives) for the revised dataset. In the worst of the 50 trials, search in the original dataset underperformed by 50% compared to what is expected by random chance), whereas the worst random trial within the final dataset still outperformed random chance by two-fold. Thus, the improved average performance and lower variability in our revised results suggests improved coherency in our revised analysis over the original published results.

**Figure 9:**
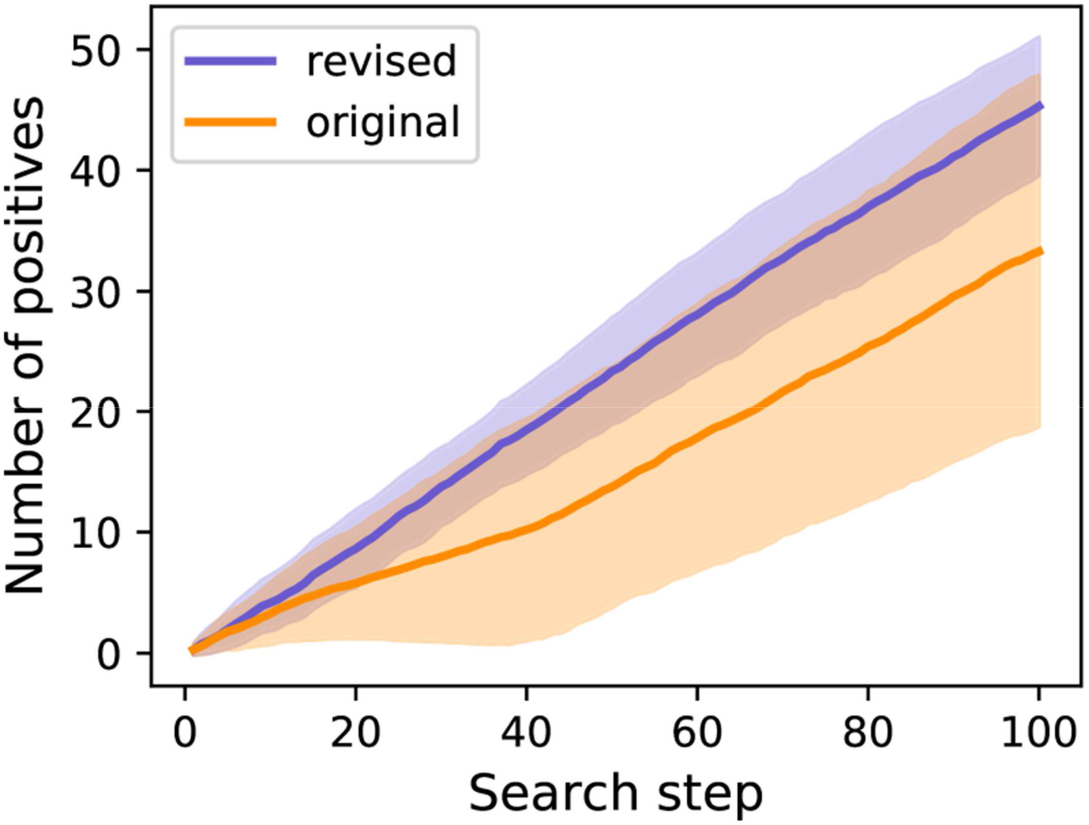
Enhanced Nonmyopic Active Search (ENS) Results. Performance of the active search algorithm ENS within each dataset (original or revised). The line represents mean result, with shading captures +/− standard deviation. In this context, ENS seeks to select each successive interaction such that the total number of positive interactions discovered is maximized.

## Discussion

Here, we performed a revised analysis of raw data from SH2 domain affinity experiments. We presented a new analysis framework which improved on the model fitting and evaluation methods of previous work. We report high-confidence true positive interactions, added thousands of true negative interactions, and removed false negative results due to inactive protein. We also report the minimum replicate measurement instead of the mean – an appropriate approach when protein concentration overestimation errors are the largest source of variance in the data.

Although raw data from only two experiments was available for detailed analysis, we were fortunate that it consisted of a large quantity of measurements from FP, a well-established solution-based experimental system commonly used for analytical biochemical assays. All in vitro experimental methods have limitations when attempting to understand behavior in vivo, but early high-throughput experiments used arrays that had limitations and biases for higher affinity interactions (*13*). Those experiments had either the peptide (*11*, *12*) or the protein (*7*–*10*) mounted on a surface, and are less preferable to a method where both molecules were measured in solution. So despite limited availability of raw data, the data available is likely to be the best type for further analysis.

We saw very high variance in affinity within replicate measurements in this data. On its face, such high variation suggests a significant problem with either experimental design or experimental method. At a minimum, it suggests that another (uncontrolled for) variable is being measured instead of the desired variable being tested. In the worst case, the remedy requires identifying and controlling for the source of variation, and redoing the experimental measurements. Even the authors of the original publication argued that the “greatest source of variability in the FP assay…is batch-specific differences in protein functionality.” (*13*) However, we have shown that the patterns found in the data are consistent with protein concentration errors, and that the likely sources of error (purification and degradation) result in overestimation of protein concentration. Because these types of errors all result in unknown amounts of active protein concentration overestimation, reporting the minimum replicate K_d_ for each domain peptide pair represents the value closest to the true activity of the protein.

In this analysis we implemented a simple metric of quality, SNR, which weights the total fit error by the size of the signal. The SNR metric was effective at eliminating suspect fits, while rejecting very few high quality measurements (Fig. S9). Nevertheless, this metric may not be appropriate for all types of data, particularly data with a large prevalence of single point outliers. We extensively explored using alternate metrics, including confidence intervals (CI). Bootstrapped CIs, established by parametric boot strapping via residual resampling, can add more information than a single fit result because they provide a range of certainty for a given measurement. However, we found that this method had significant limitations on this data, and performed worse than the SNR metric. In this data, bootstrapped CIs have even greater vulnerability to errors from outliers, are limited by small sample sizes (only 12 residuals per measurement), and suffer from heteroscedasticity of residuals (causing the high variability in low-concentration data points to be assigned to high-concentration data points) ultimately resulting in unrealistic intervals for affinity.

Several analysis methods implemented in the original publications served as sources of randomizing error, and may suggest a reason for the failure to agree with other published SH2 interaction experiments. First, background subtraction caused an unpredictable increase or decrease of affinity due to forced errors in model fitting. The magnitude of the error depended on whether the published background was higher or lower than optimum, and on the affinity of the interaction being measured. Even small deviations could result in significant errors. Another seemingly innocuous choice – averaging multiple replicates containing degraded protein – is likely to be a significant source of error in the originally published results from this experiment. Taking the mean of multiple replicates is a standard practice, but serves to randomize reported values when protein concentration overestimation is the primary source of variation.

Other high-throughput SH2 domain-peptide experiments share many critical methods with the data reviewed here. In all published experiments measuring affinity, protein was minimally filtered after production. The limited purification is likely to result in errors in protein concentration measurements due to inactive protein contaminants. Furthermore, in none of the experiments was protein assessed for activity before being measured. This has two critical consequences: the inability to separate non-binding results from negative interactions due to non-functional protein, and additional errors in active protein concentration with respect the measured protein concentration. Even if protein concentration errors were solely due to purification, it could be the cause of the significant discrepancies between published numerical results. Furthermore, incorrect use of statistical methods to evaluate models was common to all published work – particularly the improper use of the coefficient of determination (r^2^) to determine the quality of fit of a non-linear model, and using only a single model to fit data. These choices result in a high false negative rate, and mask the high variance in replicates that our revised analysis revealed. Our results suggest that, if the raw data were available, some of these issues could be corrected in other experiments. However, due to the lack of correlation between any published high-throughput SH2 domain data, and the likelihood that similar issues plague all similar data sets, we would recommend against use of these previously published data sets in future research or models of SH2 domain behavior. We further recommend that all derivative work should be carefully reviewed for accuracy.

We want to address the best uses of the revised affinity results we present, as well as the limits of the current analysis. The negative interactions we report represent a significant improvement over theoretical methods of simulating negative interactions (*18*), as they are based on real measurements rather than statistical assumptions. Furthermore, the negative interactions we report are controlled for false negative results from non-functional protein – something no other SH2 domain data can claim. Thus, our revised results have significant potential to improve the quality of models built on categorical (binary) binding data. The limitation of the quantitative data we report is that the highest affinity measured value may not be the true affinity if a fully functional protein was never measured. Nevertheless, the highest measured affinity still represents the measured value closest to the true value. However, not all variation in the data is consistent with our hypothesis of protein concentration error, and some variation may represent other unknown sources of variation which we have not controlled for. For example, one key assumption of the receptor occupancy model requires measuring the reaction at equilibrium. Since no data is provided to prove that the 20-minute incubation time given to all samples was sufficient to bring all reactions to equilibrium, it is possible that some variation is due to measurements made in non-equilibrium conditions.

It is concerning that an entire body of published work has developed from this class of problematic results. These experiments have had a wide-reaching effect in many areas of SH2 domain research: the data has been used to draw specific conclusions about SH2 domain biology such as identification of EGFR recruitment targets (*34*), to explain quantitative differences in RTK signaling (*9*), and as evidence to understand the promiscuity of EGFR tail binding (*35*). In addition, this work has been used to guide experimental design by filtering potential binding proteins by affinity (*36*), to reconcile confusing experimental results (*37*), and to guide new experimental hypothesis testing (*38*). It has played a role in cancer research as context to understand kinase dependencies in cancer (*39*), and as evidence of HER3 and PI3K connections as relevant to PTEN loss in cancer (*40*). It has influenced evolutionary analysis (*41*), been used to design mechanistic EFGR models (*42*, *43*), and has been used in algorithms for domain binding predictions (*14*–*18*, *44*).

Finally, we would like to discuss best practices for future data gathering and reporting. HTP studies have great value, and provide a vast quantity of often never before measured data. These methods have been useful to a wide variety of domain-motif interactions, for example SH3-polyproline interactions (*45*, *46*), PDZ domains interacting with C-terminal tails (*47*–*49*), and major histocompatibility complete (MHC) interactions with peptides (*50*, *51*). However, just as quickly, errors in these studies propagate rapidly and thereby into research results of other investigators. This suggests that an even higher than normal standard of care is necessary when evaluating such publications. A set of best practices for HTP methods should be established in the community. We recommend all raw data from high throughput experiments should be published, along with all code used to process that data. This would make the initial data far more valuable for future research, much like the raw arrays stored Gene Expression Omnibus (GEO), or the raw experimental measurements are stored along with the protein structure in the Protein Data Bank (PDB). To this end, we have provided the original raw data and our full revised data (including intermediate steps) on Figshare (DOI: https://doi.org/10.6084/m9.figshare.11482686.v1), and provided the code for the analysis pipeline on GitHub (https://github.com/NaegleLab/SH2fp) so that future evaluation can be more easily accomplished by other researchers. Although portions of our code are highly specific to the format of these datasets, the code is written in a modular fashion that can be easily repurposed in other studies. We also recommend that methods for quantifying activity should be a best practice in studies quantitatively measuring protein. Alternatively, methods which do not depend so heavily on accurate protein concentration should be preferred. One such concentration-independent method of measuring interaction affinity was recently developed by the Stormo lab (*52*). In that method, a 2-color competitive fluorescence anisotropy assay measures the relative affinity of two interactions in solution. By measuring interaction against two peptides at once from the same pool of proteins, the concentration of the protein and the proportion of active protein is the same in both interactions. When the ratios are calculated, the concentration and activity drop from the calculation of affinity. Although this method only provides relative affinity, if one could carefully establish absolutely affinity for a single peptide (or panel of peptides), absolute affinity could be extended to all interactions. Another recent experiment also uses competitive fluorescence anisotropy, but measures a competitive titration curve in a single well with an agarose gradient (*53*). Diffusion forms a spatiotemporal gradient for the interaction, and so one can produce a full titration curve in each well in a multi-well plate, measuring both affinity and active protein concentration simultaneously. Regardless of the specific method, it should be a best practice to account for or control for the concentration of active protein within the measurement of total protein concentration.

## Methods

### Raw Data

Upon receipt of the Jones 2012-14 raw data, we examined the data for consistency and completeness. We found that the data did not cover all interactions described in the original publication. However, by limiting our revised analysis to interactions of single SH2 domains with phosphopeptides from the ErbB family, as well as KIT, MET, and GAB1, we were able to limit the effect of missing raw data. Within this scope, only a handful of individual replicate interactions were then missing (approximately 138 replicate-level measurements out of over 37,000 measurements) and were limited to 3 domain-peptide pairs. Fortunately, two of the domain-peptide pairs were represented by other replicate measurements. The data we examined for this revised analysis cover the interactions of 84 SH2 domains with 184 phosphopeptides. The peptides came from receptor proteins from the four ErbB domains (EGFR/ErbB1, HER2/ErbB2, ErbB3, ErbB4) as well as KIT, MET, and GAB1. Of SH2 proteins containing a single SH2 domain, 66 domains were measured: ABL1, ABL2, BCAR3, BLK, BLNK, BMX, BTK, CRK, CRKL, DAPP1, FER, FES, FGR, GRAP2, GRB2, GRB7, GRB10, GRB14, HCK, HSH2D, INPPL1, ITK, LCK, LCP2, LYN, MATK, NCK1, NCK2, PTK6, SH2B1, SH2B2, SH2B3, SH2D1A, SH2D1B, SH2D2A, SH2D3A, SH2D3C, SH3BP2, SHB, SHC1, SHC2, SHC3, SHC4, SHD, SHE, SHF, SLA, SLA2, SOCS1, SOCS2, SOCS3, SOCS5, SOCS6, SRC, STAP1, SUPT6H, TEC, TENC1, TNS1, TNS3, TNS4, TXK, VAV1, VAV2, VAV3, and YES1. From SH2 proteins with double domains, C-terminal and N-terminal domains were individually measured from 10 proteins: PIK3R1, PIK3R2, PIK3R3, PLCG1, PTPN11, RASA1, SYK, ZAP70, PLCG2 (N-terminal only) and PTPN6 (C-terminal only). One peptide had no measurements in the raw data (EGFR pY944). Within this revised scope, the available raw data covered approximately 99.6% of the originally available raw data.

The raw data for each measured interaction consisted of fluorescence polarization measurements of an SH2 domain in solution with a phosphopeptide at 12 concentrations. The measurements were arranged on 384 well plates: 32 different SH2 domains at each of 12 concentrations, all measured against a single peptide per plate. Protein concentrations represented 12 serial dilutions of 50% starting with either 10 μM or 5 μM protein.

### Model Fitting, Model Selection, and Replicate-Level Calls

For each replicate measurement, we fit two models: the linear model (equation 4) and the receptor occupancy model (equation 2). Model fits were evaluated with the bias corrected Akaike Information Criterion (AICc), and the model with the lower AICc score was selected (*19*).

The Akaike Information Criterion (AIC) as a quality metric, was calculated by

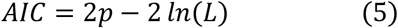

where *p* is the number of parameters in the model, and *ln(L)* is the maximum log-likelihood of the model. In a non-linear fit, with normally distributed errors, *ln(L)* is calculated by

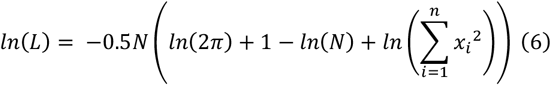

where x_1_, …, x_n_ are the residuals from the nonlinear least squares fit and N is the number of residuals. The bias corrected form of AIC, referred to as AICc, is a variant which corrects for small sample sizes, e.g. when one has fewer than 30 data points. AICc is calculated as follows:

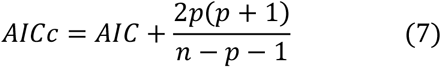

where n is the sample size, and p is the number of parameters in the model (*19*). Each replicate had a sample size of 12. The receptor occupancy model had three parameters (affinity (K_d_), saturation level (F_max_), and offset (F_0_)), while the linear model had two parameters (slope (*m*), and background offset (F_0_)).

Replicates that were fit best by the linear model with a slope of less than or equal to 5mP/μM were categorized as negative interactions, or ‘non-binders’. Linear fits with a slope greater than 5mP/μM were categorized as aggregators. Replicates that were fit best by the receptor occupancy model were subsequently evaluated for signal to noise ratio (SNR, equation 3). If the SNR was greater than one, the replicate was categorized as a positive interaction or ‘binder’, otherwise, it was rejected as a low-SNR fit and removed from consideration.

### Identifying Non-Functional Protein

Once all individual fits were complete, runs were examined for non-functional protein. If an entire run lacked even one positive binding interaction, and those same interactions measured positive on another run, the non-binder, aggregator, and low-SNR calls on that run were changed to non-functional protein and removed from consideration.

### Replicate Handling for Domain-Peptide Measurements

For each domain-peptide pair, only replicates that were marked as binders with sufficiently high signal to noise ratio (SNR) were considered. For a given domain-peptide pair, the minimum numeric value of K_d_ (representing the strongest affinity) was reported as the final K_d_ for that domain peptide pair.

### Active search

The probability model (*33*) used a simple k-nearest neighbor (k = 20) where distance is defined by average Euclidean distance of corresponding divided physicochemical property scores (DPPS) features of the amino acids (*54*) comprising the peptide, i.e.:

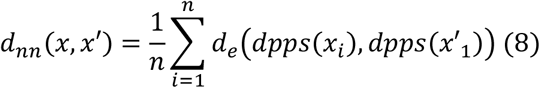

where *d*_*nn*_ is the distance used to define nearest neighbors, *d*_*e*_ is the Euclidean distance, *n* is the number of amino acids in the peptide (here *n* = *9*), and *dpps(x*_*i*_) is the *DPPS* feature vector of the i^th^ amino acid in peptide *x*.

## Acknowledgements

We would like to thank Richard Jones, Ron Hause and Ken Leung for providing the raw data required for this analysis.

## Supporting Information

Final Revised Affinity Data.xlsx – contains the interaction affinities between domains and phosphopeptides based on our revised analysis.

**Fig. S1:**
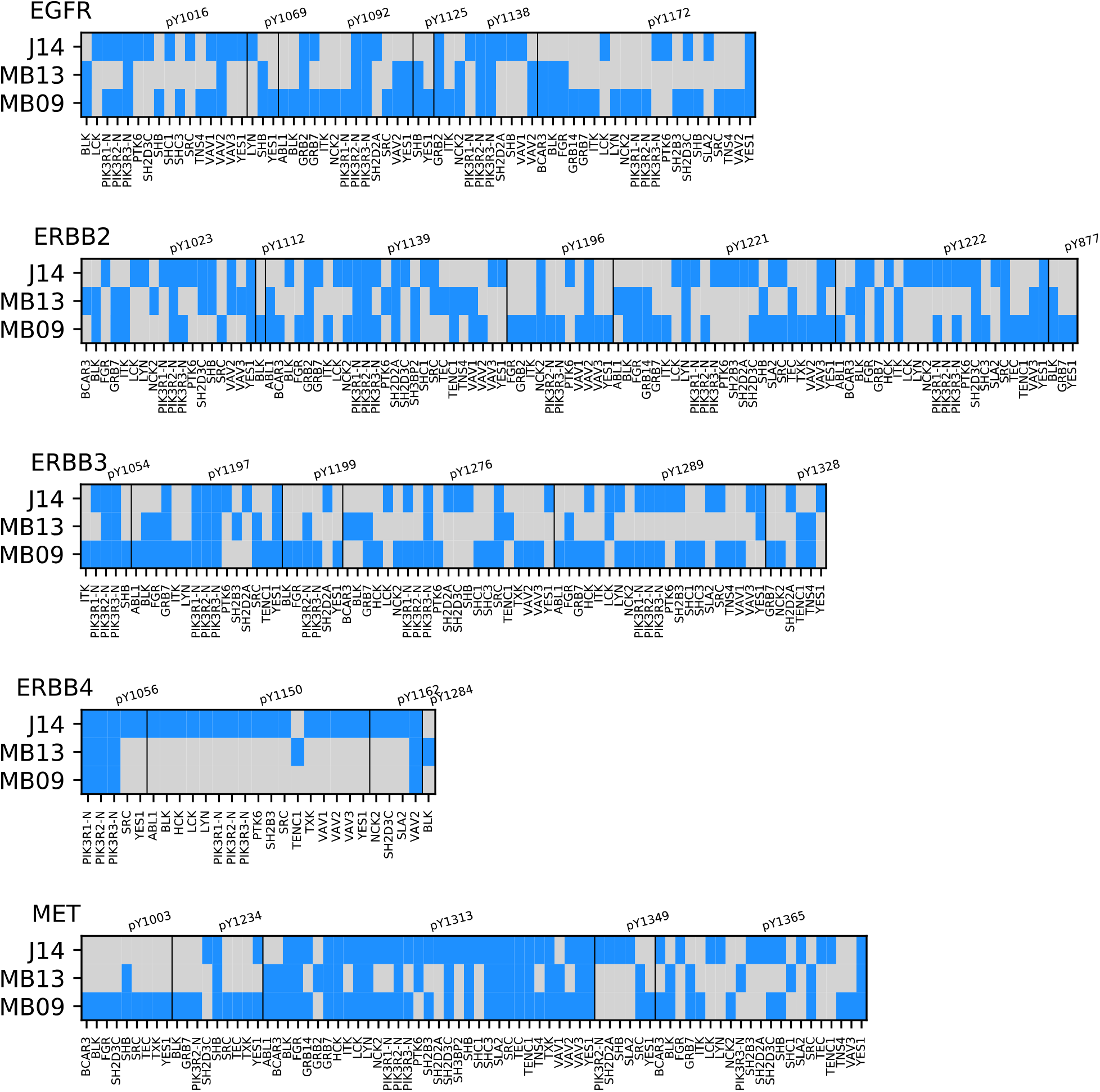
Domain-Peptide-Level Comparison of Binding Between Published Results. Blue denotes positive interations (binding). J14: Jones 2012-14; MB13: MacBeath 2013; MB09: MacBeath 2006-09.

**Fig. S2:**
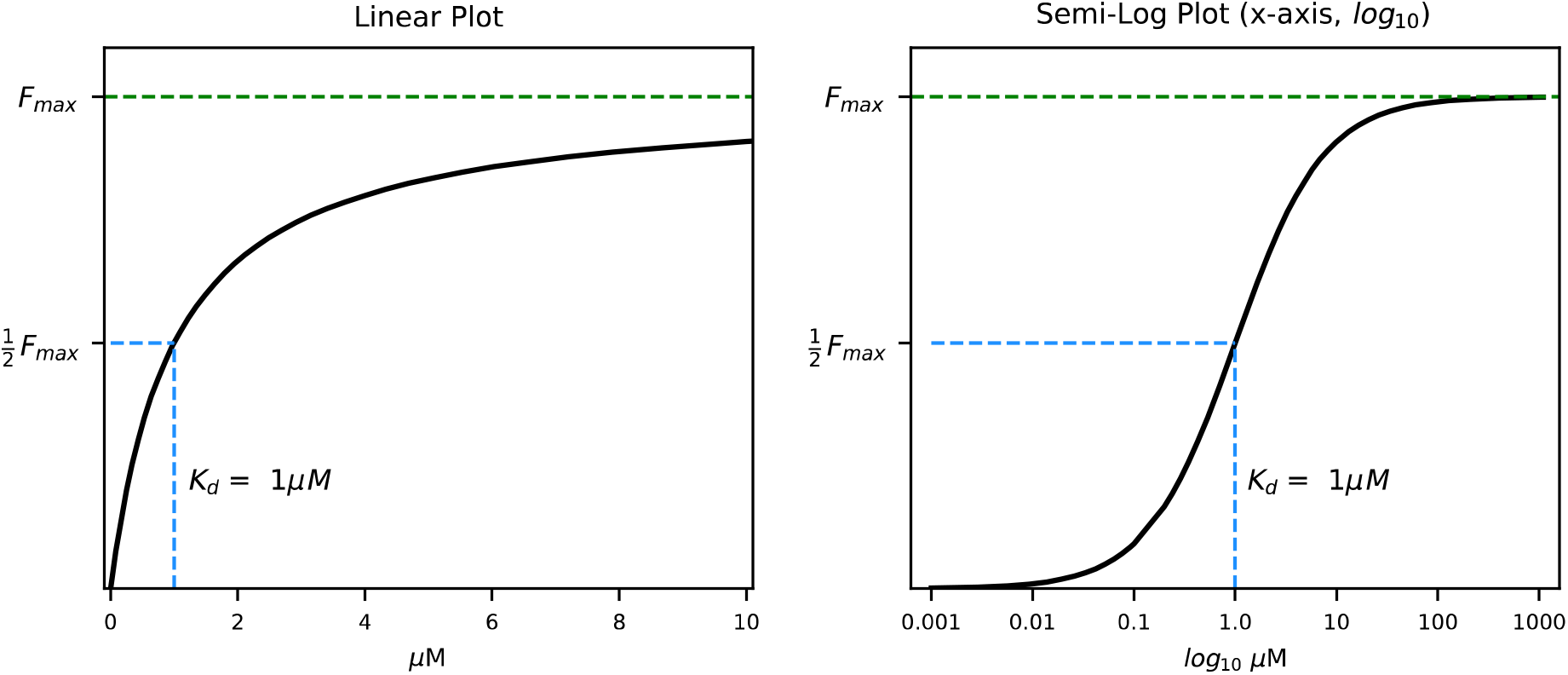
The Receptor Occupation Model. Saturation plots in both linear (left) and semi-log (right) demonstrate increasing saturation with increasing fractional occupancy, with full saturation achieved at F_max_. The affinity (quantified by the K_d_) can be derived by fitting a curve to the data, but can also be derived graphically as seen above. At equilibirum, the K_d_ is equal to the concentration when ½ of the receptor is occupied (½ F_max_). In the semi-log curve, where the concentration axis is in log_10_ scale, K_d_ can be identified easily because it corresponds with the inflection point at the center of the the s-shaped curve.

**Fig. S3:**
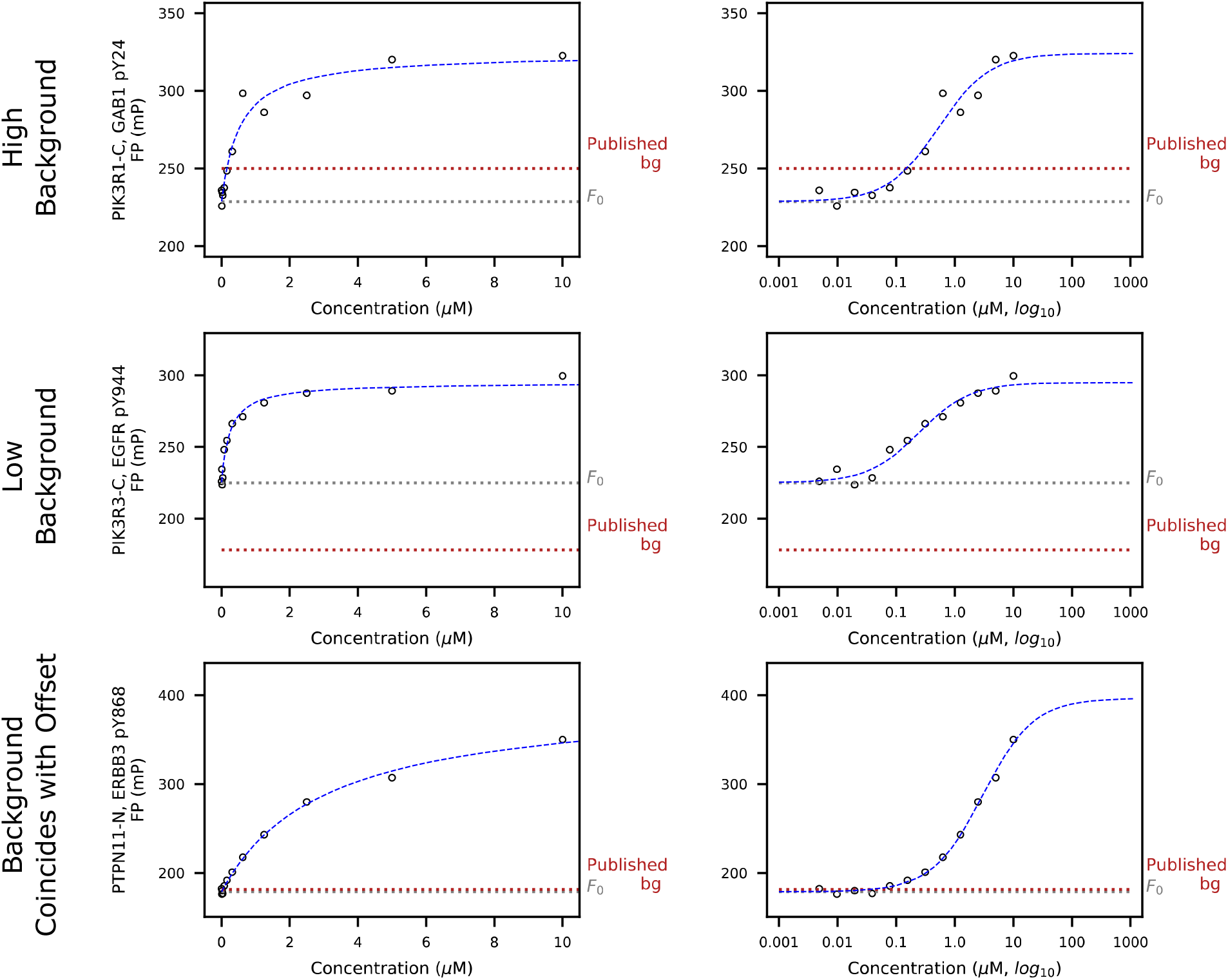
Examples of Background Fluorescence. The raw data exhibited highly varying levels of background FP. The background FP level should represent the level below which measurements are random noise. For some measurements, background values were higher than many of the individual low-concentration measurements (first row). In these cases, the high quality of the data and the fits seem to contradict the limits imposed by the reported background. In other cases, the background values were significantly lower than the signal level of the lowest concentration measurements (second row). In some cases, the published background value matched the offset value from the model fit (third row). Because of the discrepancy with background values we found and the use of the background subtraction method in the original publication, we decided to evaluate the background data and quantify the effect of background subtraction on the accuracy of the model fits (see Fig. S4).

**Fig. S4:**
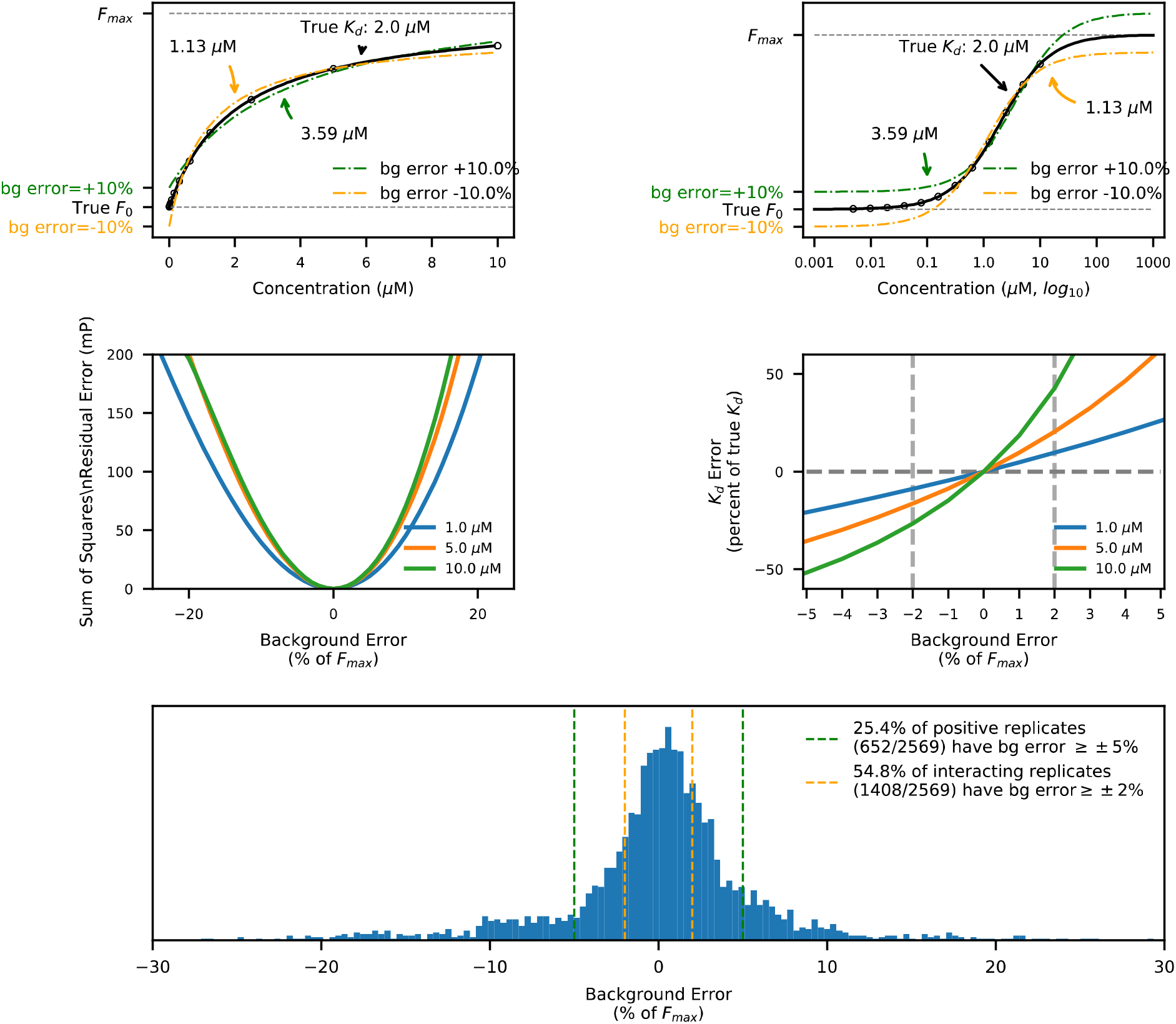
Background Subtraction Causes Errors in Receptor Occupancy Model Fits. The background subtraction method forces the receptor occupancy model curve through a fixed y-axis point (the background level) at zero concentration (top row; most easily seen on the semi-log plots, top row right). For example, given a theoretically perfect measurement with a K_d_ of 2.0μM (top row, black line), background subtraction induces an error in affinity. A background 10% higher than optimal (top row, green) causes a change in the fitted curve (top row, green dashed line), which results in a derived K_d_ of 3.59μM (a +179.5% error). A background 10% lower than optimal (top row, orange) causes a change in the fitted curve (top row, orange dashed line), which results in a derived K_d_ of 1.13μM (a −40.5% error). Background subtraction always results in an increase in residual error (middle row left), which results in an error in the fitted affinity parameter (middle row, right). Background errors are nonlinear with affinity, and result in larger errors at lower affinities (middle row, left and right). A background error of ±5% results in a −21%/+26% error in K_d_ at 1.0μM, and a −52%/+195% error in K_d_ at 10.0μM. Even a 2% background error results in a −9%/+10% error in K_d_ at 1.0μM, and a −27%/+43% error in K_d_ at 10.0μM. Over 25% (652/2569) of all replicate measurements that demonstrate positive interactions have background errors greater than ±5%, and over 54% (1408/2569) have background errors greater than ±2%. Thus, background subtraction causes errors in affinity which can increase or decrease affinity based on whether the background is high or low, and are non-linear by affinity. Since the relationship of the background is seemingly random, and the error factors are non-linear, background subtraction acts as a significant source of random error. Based on these findings, we rejected the background subtraction method in favor of a fitted offset (Equation 2). Background Error is expressed as a percent of the saturation value, and is defined as the difference between background and True F_0_ divided by the difference between True F_0_ and F_max_ (saturation).

**Fig. S5:**
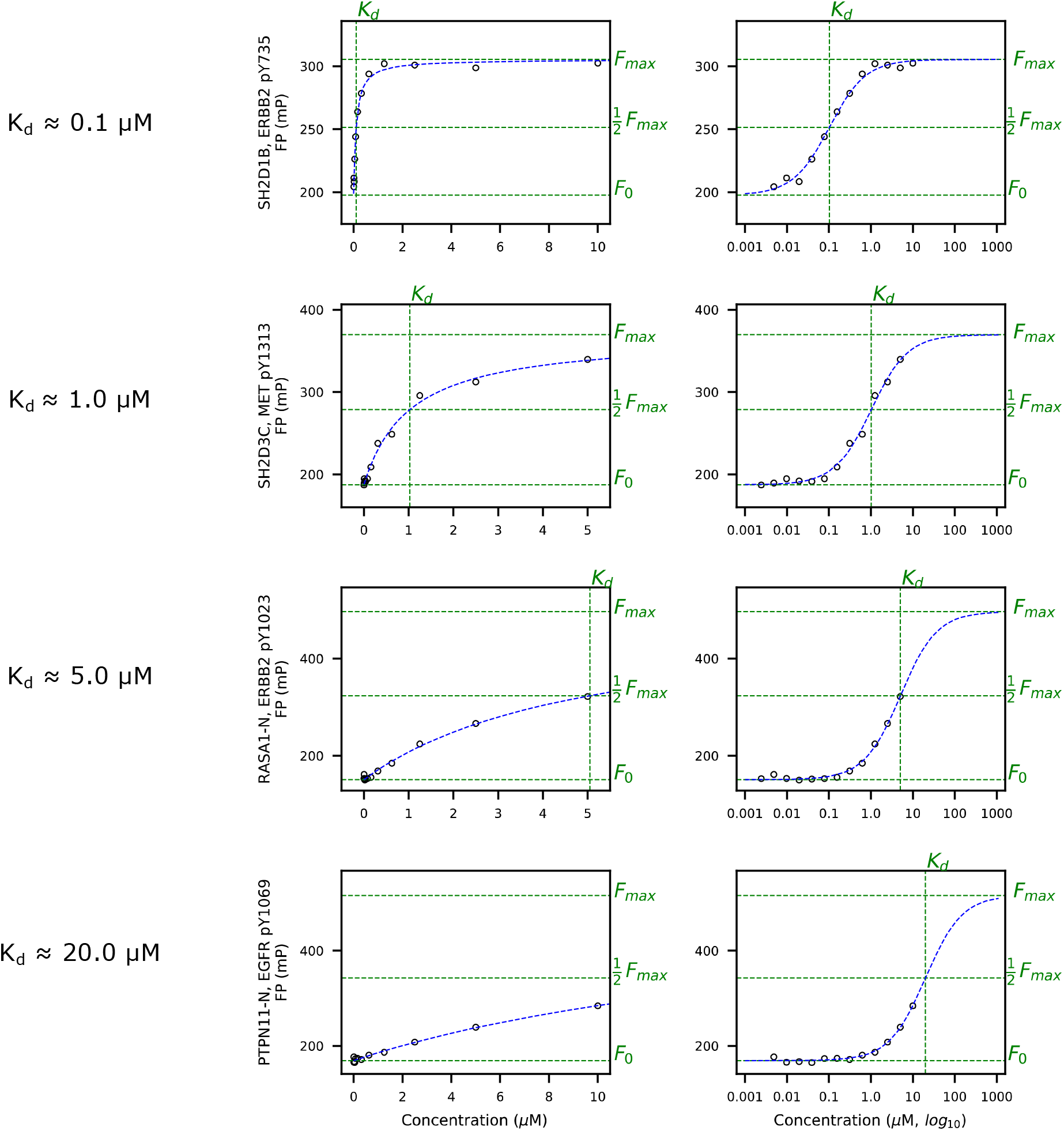
Receptor Occupancy Model Fits for Various Affinity Interactions. High-quality receptor occupancy model fits showing positive interactions at varying affinities (K_d_) from 0.1μM to 20μM. The range of concentrations spanned by each 12-concentration measurement was either 2.4nM–5μM or 4.9nM–10μM. For an ideal binding saturation experiment attempting to quantify K_d_, the concentrations tested should span above and below K_d_, and the highest and lowest measured concentrations should establish the plateaus seen on semi-log saturation plots (all rows, second column). For interactions with 0.1 μM K_d_ (first row), on the semi-log plots, data points are evenly distributed on either side of the inflection point, and establish the lower plateau of no signal and upper plateau of saturation. For interactions with a K_d_ of 1.0 μM (second row), the upper plateau of the semi-log saturation curve no longer has any coverage from the data. For interactions with K_d_ > 5μM (rows 3 and 4), no data points are found above Kd, which significantly increases potential inaccuracies in model fitting. Thus, the concentration ranges chosen make this experiment best suited to identify affinity in the 0.05μM to 0.5μM range. Since data in the original publication was reported up to 20μM, results with low affinities (higher K_d_ values) are likely to be less accurate. In addition, this pattern of coverage suggests that every data point is critical for accuracy, particularly for concentrations above K_d_.

**Fig. S6:**
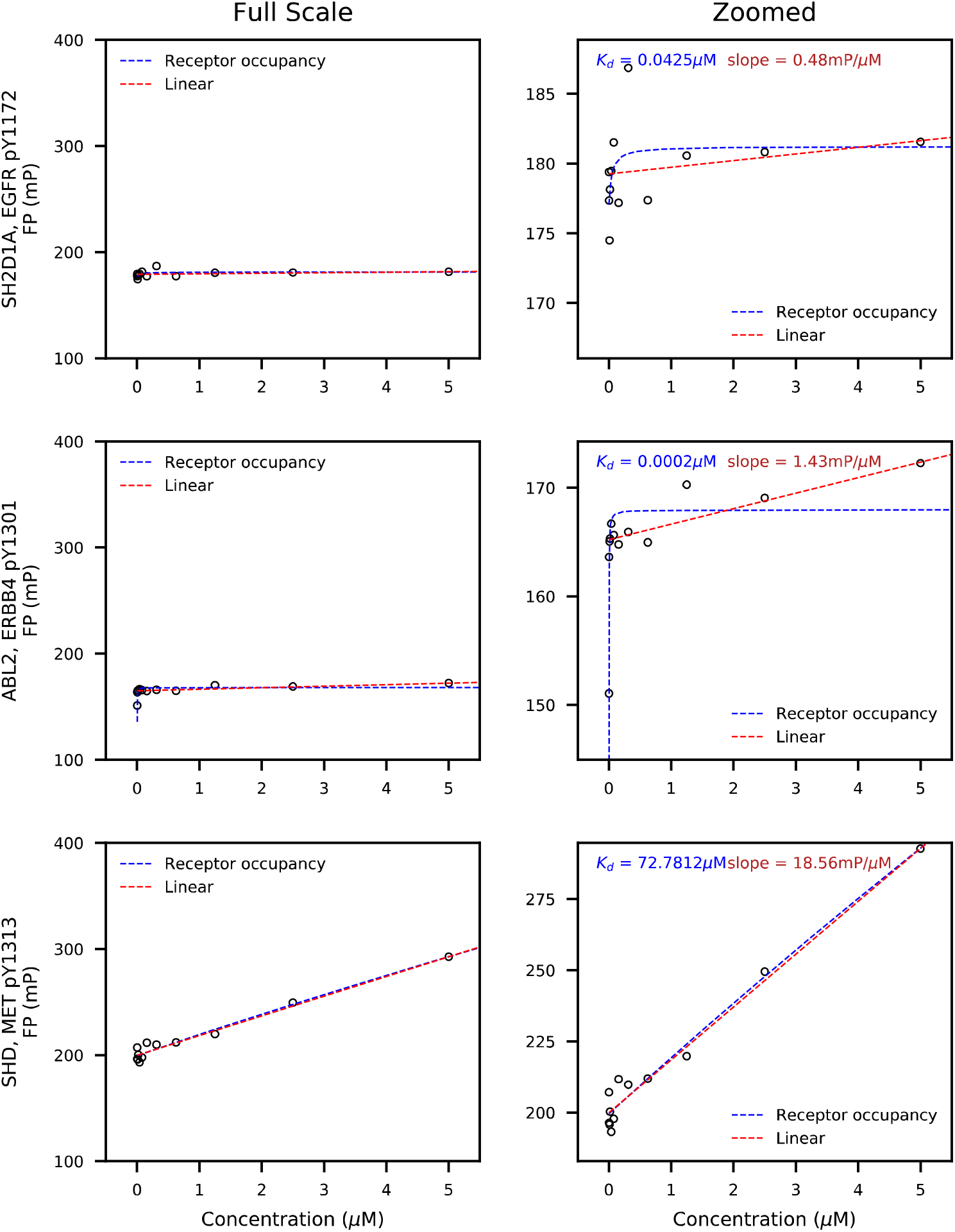
The Receptor Occupancy Model Fails to Accurately Fit Non-binding Measurements In Practice. Non-binding data generally manifests as data points in a roughly horizontal line, with a level of superimposed noise (row 1, see first column for examples at full scale). Upon close examination (row 1, see second column for zoomed view of the same measurement), noise in individual data points can be more clearly visualized. Using a least-squares algorithm to fit the Receptor Occupancy model can result in fit artifacts for non-binding data. The fitting errors follow two patterns: In the first pattern, noise in the data is over-fit, resulting in a rapidly saturating curve, rather than a straight horizontal line (row 1, blue dashed line). This saturation curve poorly fits the data and often has a low saturation value (on the order of 5mP units). Ironically, this artifact results in miscategorization as a binder with a high affinity, rather than the true result reflecting a failure to interact. In a similar type of fit-artifact (row 2), all but one data point is considered to be at saturation, while one single point sets the rest of the saturation curve, resulting in an artifically low K_d_. In the third pattern (row 3), when there is non-specific binding or aggregation present, data can also present as a line with a high slope showing no signs of saturation. A saturation value of this size cannot result from the one-to-one interaction assumption of the receptor occupancy model, and clearly represents a fit artifact, likely aggregation or other non-specific binding.

**Fig. S7:**
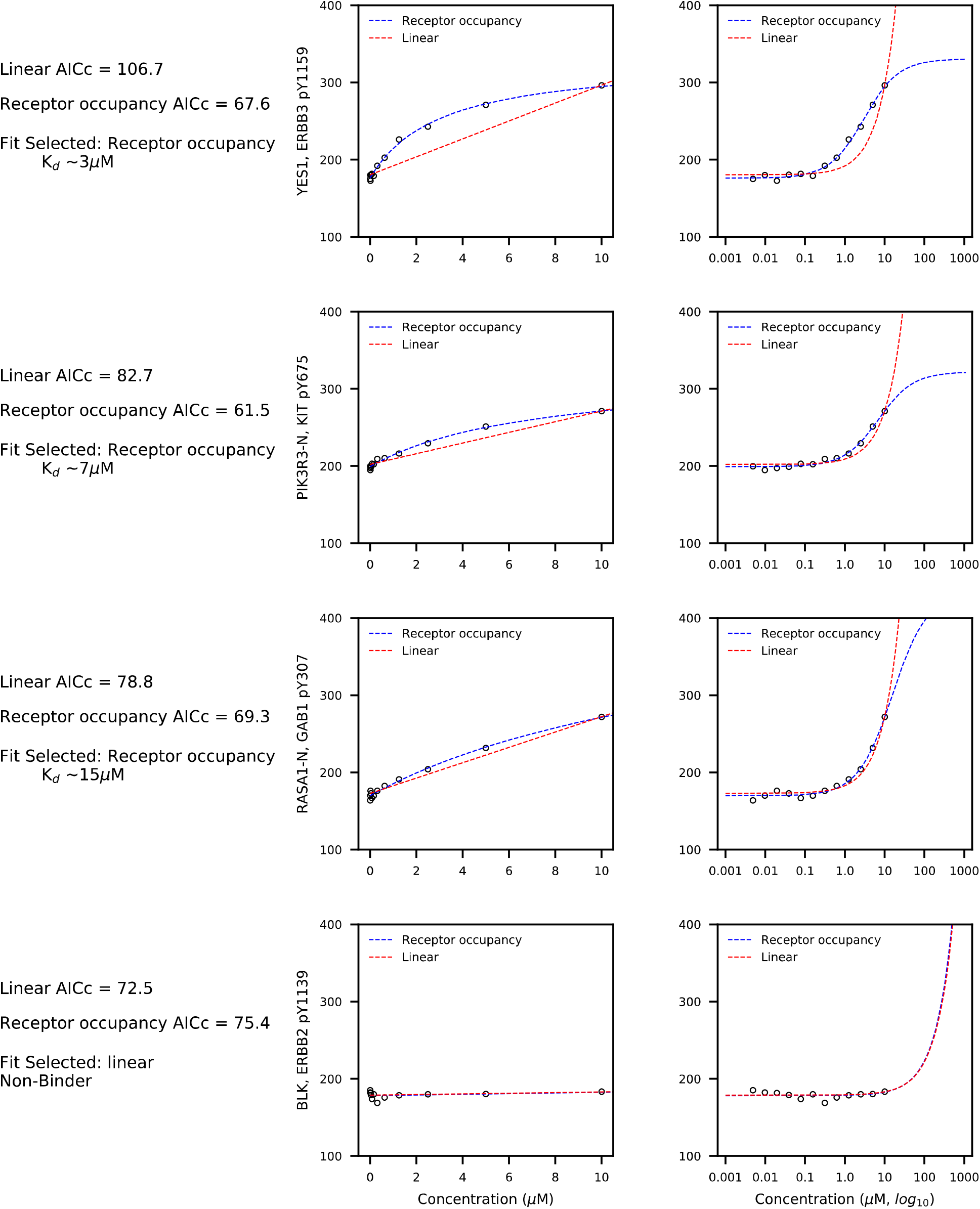
Model Selection Examples Using AICc. Both linear and receptor occupancy models are fitted to the data. AICc scores are calculated and compared between models – the model with the lowest AICc score is selected as the best fit. If a linear fit is chosen, and the slope is less than 5mP/μM, the interaction is classified as a non-binding interaction.

**Fig. S8:**
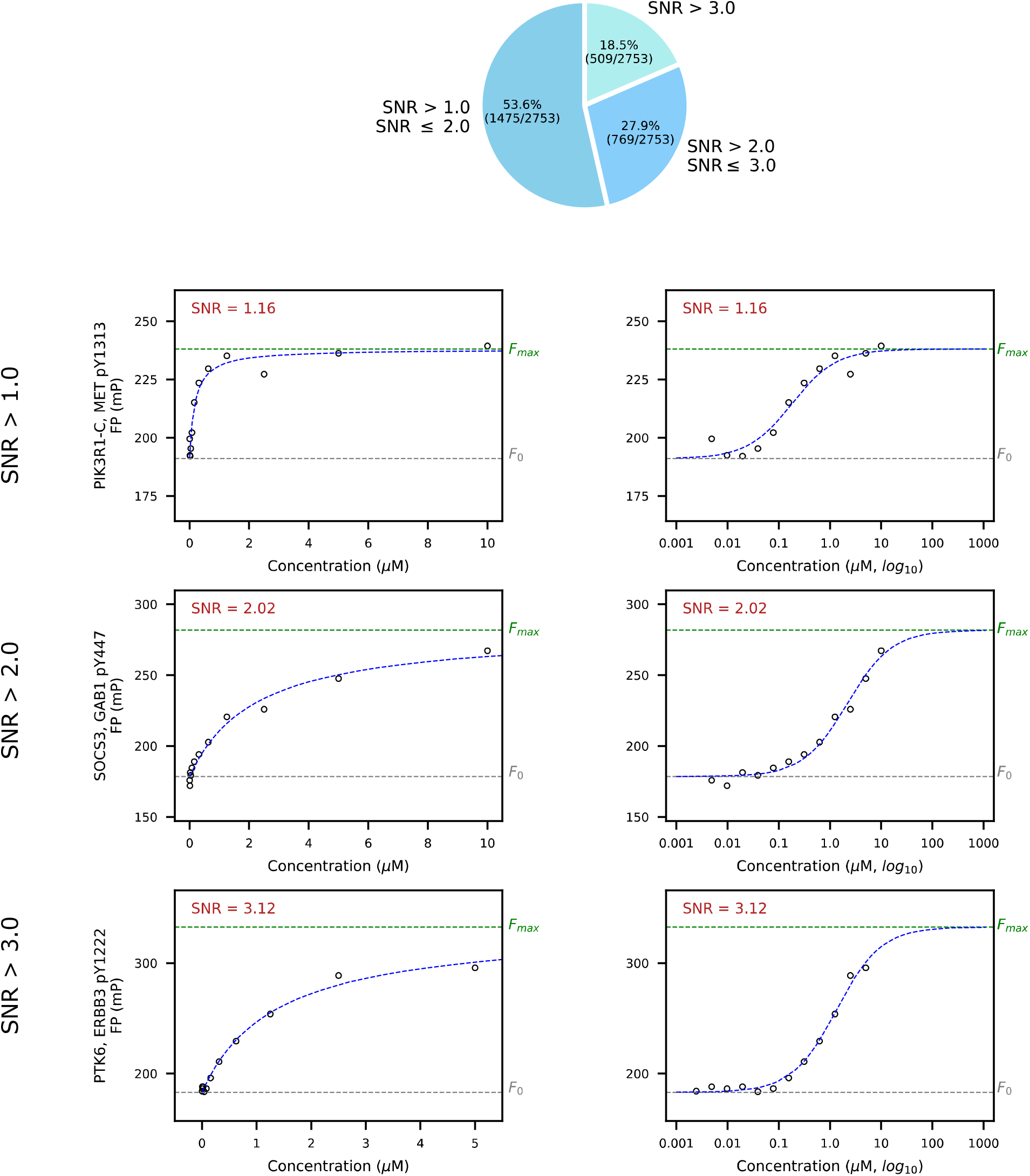
Model Fitness Examples Using The Signal-to-Noise Ratio (SNR). The model fitness metric, deemed signal-to-noise ratio (SNR), evaluates the magnitude of residual errors of fit to the model (a form of noise), and weighs this sum by the overall size of the fluorescent signal measured. As can be seen from the examples above, a signal to noise ratio of 1.0 or greater represents high-quality fits to the model, with little deviation from the model fit line.

**Fig. S9:**
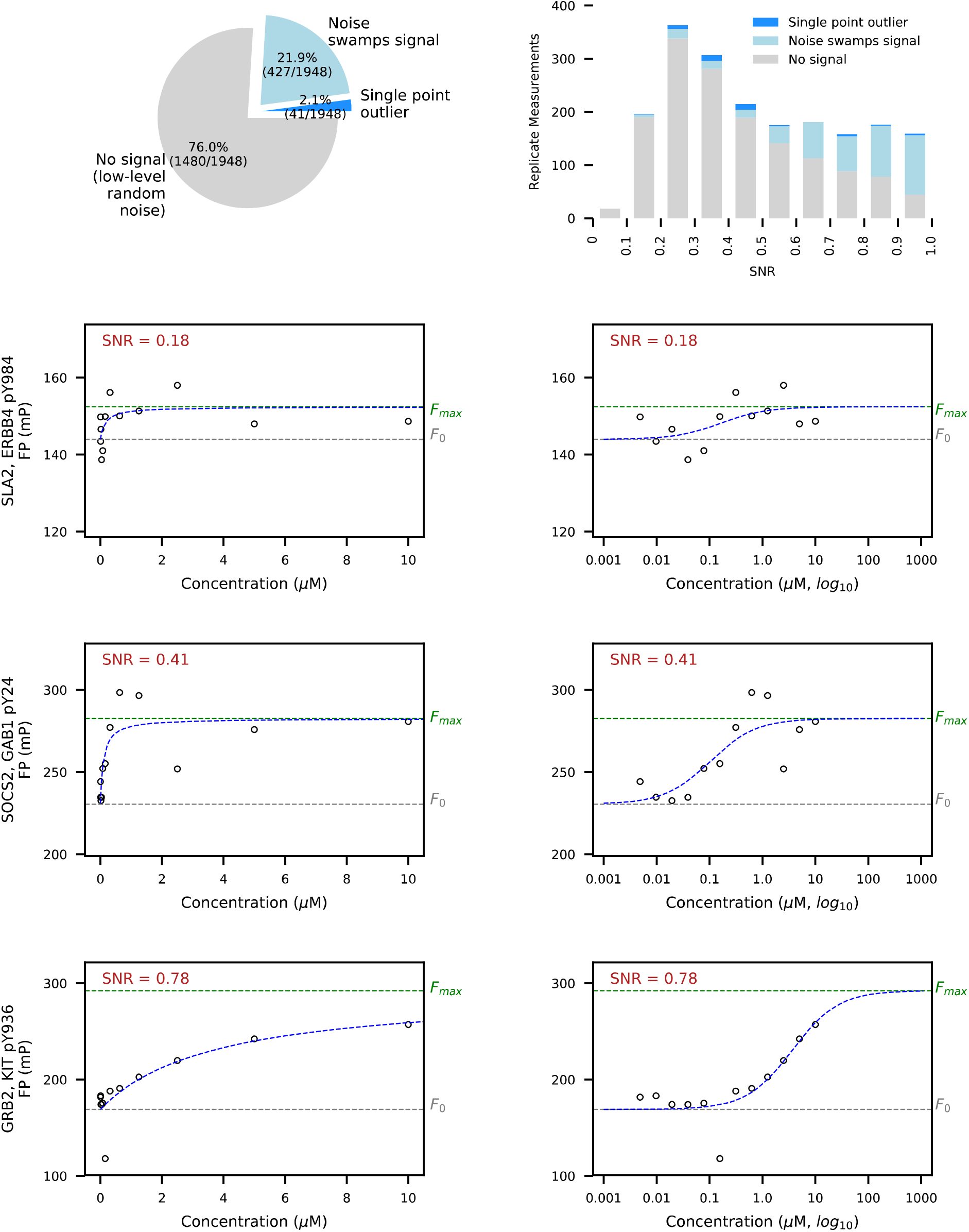
Classification and Quantification of Fits with SNR < 1.0. Replicates with SNR<1 made up 5.2% of all fits (1948/37378). These low-SNR fits fell into three classes: (1) no signal fits that nevertheless fit the receptor occupancy model better than a linear model, but which consisisted soley of low-level radom noise (76.0%, 1480/1948, row 2); fits with some non-zero signal present but so swamped by noise as to not resemble a saturation curve 21.9%, 427/1948, row 3), and otherwise good fits to the receptor occupancy model with one large outlier (2.1%, 41/1948, row 4) pulling down the SNR metric below 1. Although the metric excludes some viable measurements that would be kept when reviewing by eye (e.g. 41 single point outlier replicate measurements) this represents only 1/10th of 1% of all replicate measurements (0.11%, 14/37378). The composition of the three classes varied with SNR value (top row, right). As SNR increases, fewer non-signal measurements are encountered and more non-zero signal measurements occur. The metric predominantly functions to exclude artefactual receptor occupancy fits to random noise in zero-signal measurements. See the Discussion for thoughts on alternate metrics.

**Fig. S10:**
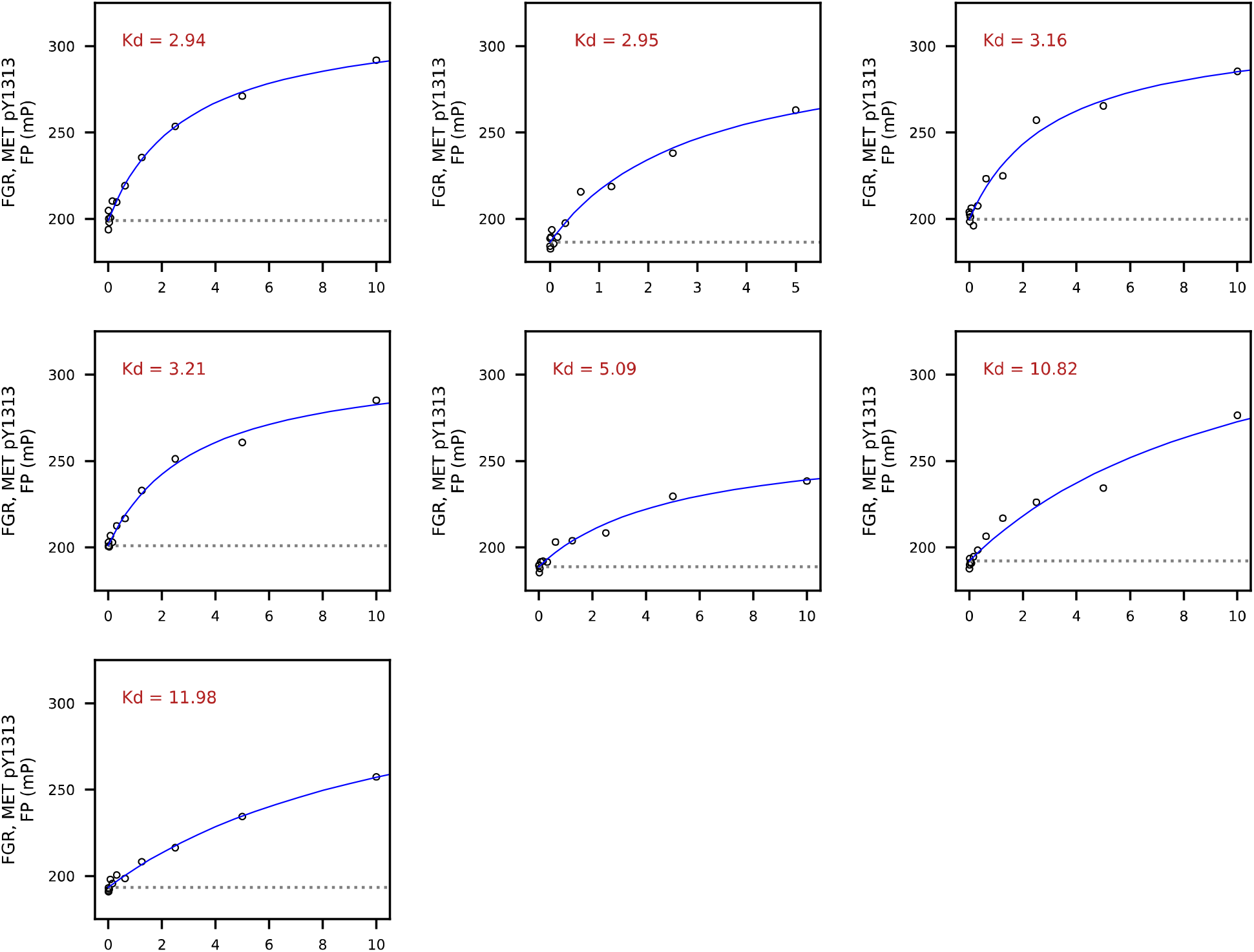
Replicate Measurements for FGR Interactions with MET pY1313. An example of high-replicate variation across replicates for a single domain-peptide pair. Each individual measurement represents a high-quality fit to the receptor occupancy model, yet the resulting affinities vary from ∼3μ m to ∼12μM. It is clear from the quality of each measurement that the variation is not due to noisy data, or fitting artifacts. Rather, each measurement seems to be a high-quality result of different affinity behavior.

**Fig. S11:**
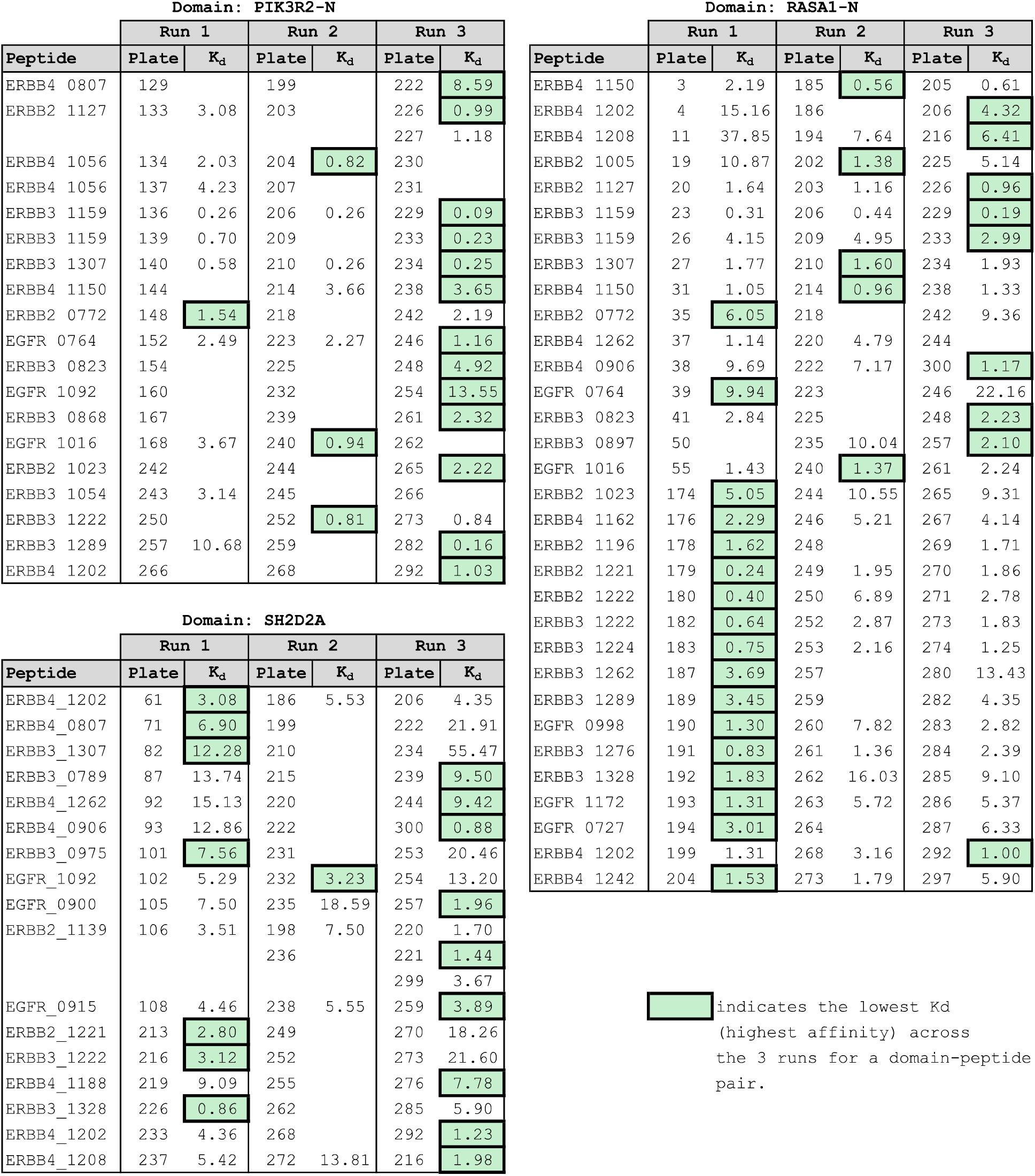
Examples of Time-Dependent Affinity Patterns in Domain Data. Variance in affinity from a systemic source can manifest as non-random patterns of variance in time. Although we don’t have an exact time for each measurement (and the same peptides were typically measured only once per run) we do have a pseudo-time substitute in the data. On each run, the peptides were measured in approximately the same order, and hundreds of peptides were measured in each run, which allows us to see patterns of protein affinity over time and across peptides from run to run. For example, in PIK3R2-N (upper left panel) we see that Run3 replicates almost always have lower K_d_ values (higher affinities) than replicates from other days. This pattern of run to run variation suggests that the protein samples tested in Runs 1 and 2 may have been degraded or from different protein batches with varying levels of impurities. For RASA1-N (right panel), no single day dominated the highest affinity until plate 174, after which the highest affinity replicates all come from Run 1. A protein sample exhausted mid-run and replaced with a fresh sample, could manifest as a surge of increased affinity in the middle of a run of lower affinity, such as seen in Run 1. Not all protein data shows such clear patterns. For SH2D2A (lower left panel), significant variation appears during Runs 1 and 3 with no run showing the lowest K_d_, but Run2 shows consistently higher K_d_ values, as well as many failures to detect interactions (blank K_d_ values) suggesting degradation or impurities on the protein from Run 2. The patterns for SH2D2A are not as clearly consistent with a simple concentration error hypothesis, and may be indicative of additional sources of variation.

**Fig. S12:**
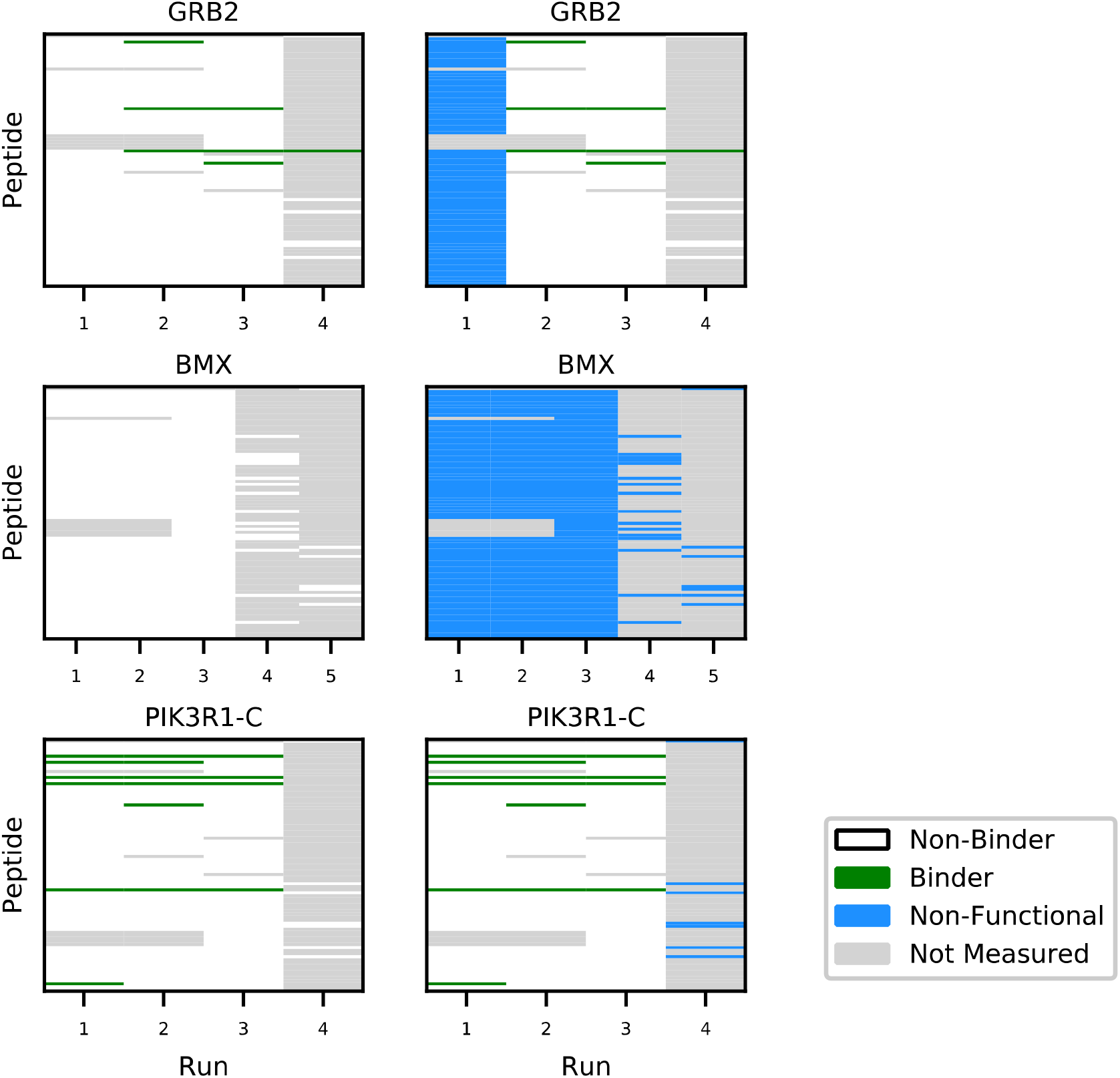
Non-Functional Protein Identification – Examples. In order to examine the data for patterns of non-functional protein, we plotted affinity by domain and by run. The results before non-functional protein identification (NFPI, left column), after NFPI (right column) are shown for three protein domains. A lack of even one positive interaction on an entire run is suggestive of non-functional protein. When other runs of the same protein show positive interactions, the runs with no positive interactions are considered to be non-functional and all measurements in these runs for the protein are removed from consideration. For example, with GRB2 (row 1), runs 2 through 4 showed some positive interactions. On run 1, however, no measurements indicated positive interactions. The lack of even one positive interaction in run 1 suggests that the protein was completely degraded or non-functional. The presence of positive interactions in the other runs acts as a positive control. Run 1 is then marked in blue in the right panel for GRB2, and removed from consideration. A different case of non-functional protein can be seen with BMX (row 2). For BMX, no positive interactions were found on any run. Although it is a formal possibility that BMX simply binds none of these peptides, we simply have no information that the protein was ever active, thus we conservatively identify all runs as non-functional. For PIK3R1-C, no measurements on the fourth run were positive interactions, while other runs contain positives, thus run 4 was categorized to be non-functional. A binder is identified by a green cell, a non-binder by a white cell, non-functional protein by a blue cell, and a non-measured interaction by a gray cell.

**Fig. S13:**
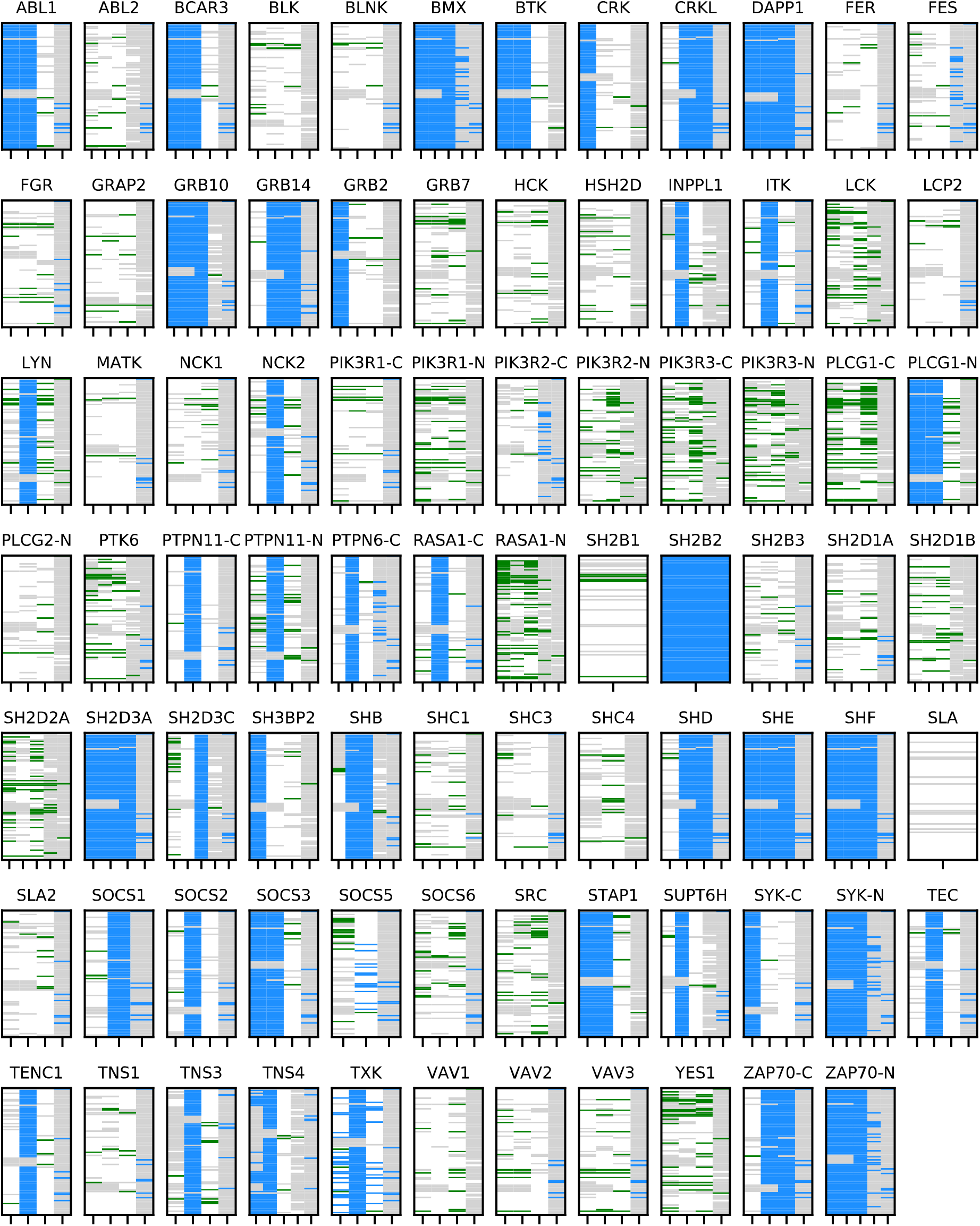
Non-functional Protein in Hause, et al (2012). Non-Functional protein results for all measured interactions from the first publication, Hause, et al (2012). See legend from Fig. S11.

**Fig. S14:**
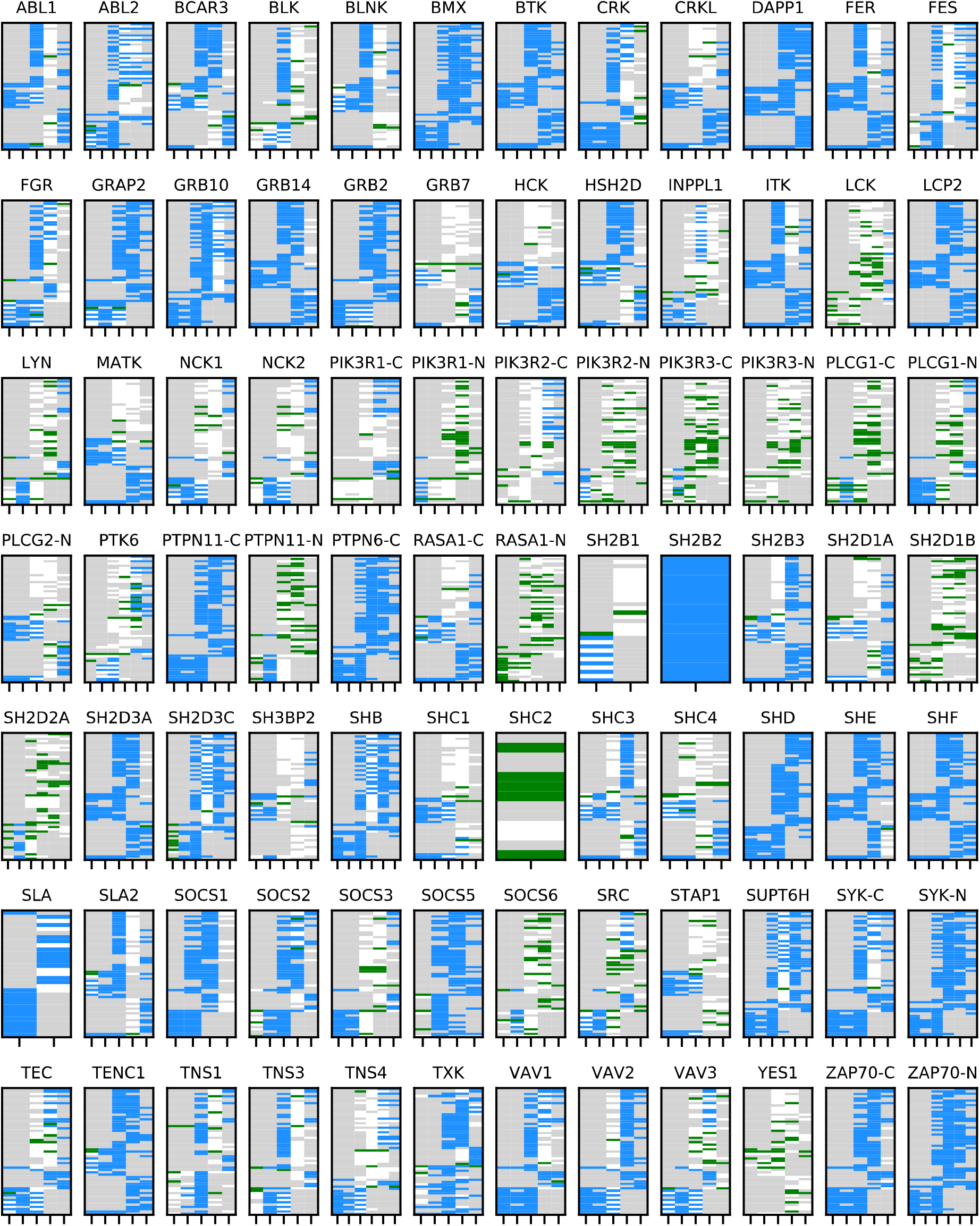
Non-functional Protein in Leung, et al (2014). Non-Functional protein results for all measured interactions from the second publication, Leung, et al (2014). See legend from Fig. S11.

**Fig. S15:**
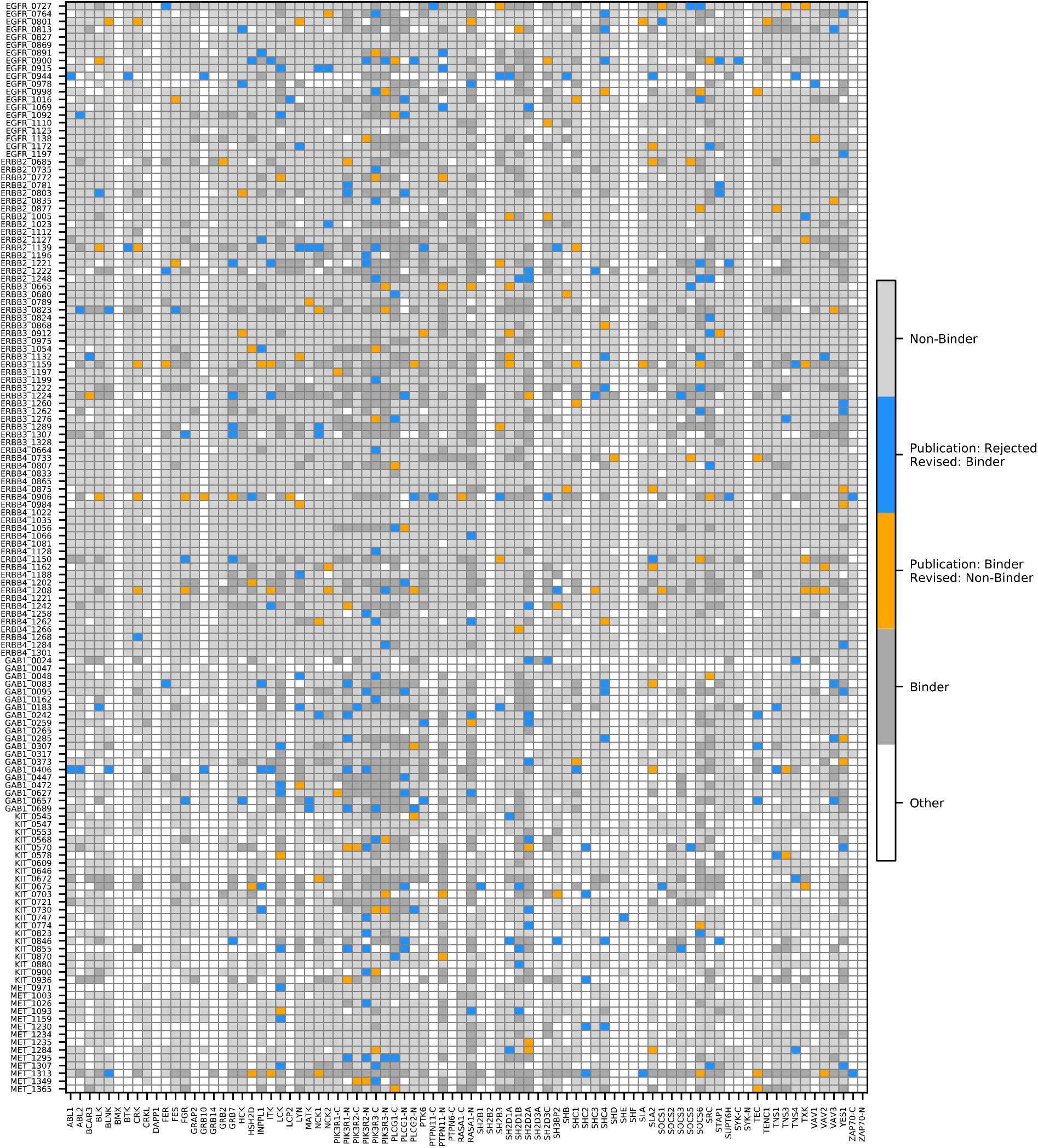
Changes In Calls Between Original Publication and Revised Analysis. A heat map showing the changes in calls in our revised analysis. Differences in calls with the original publication are found across all domains and all peptides.

**Fig. S16:**
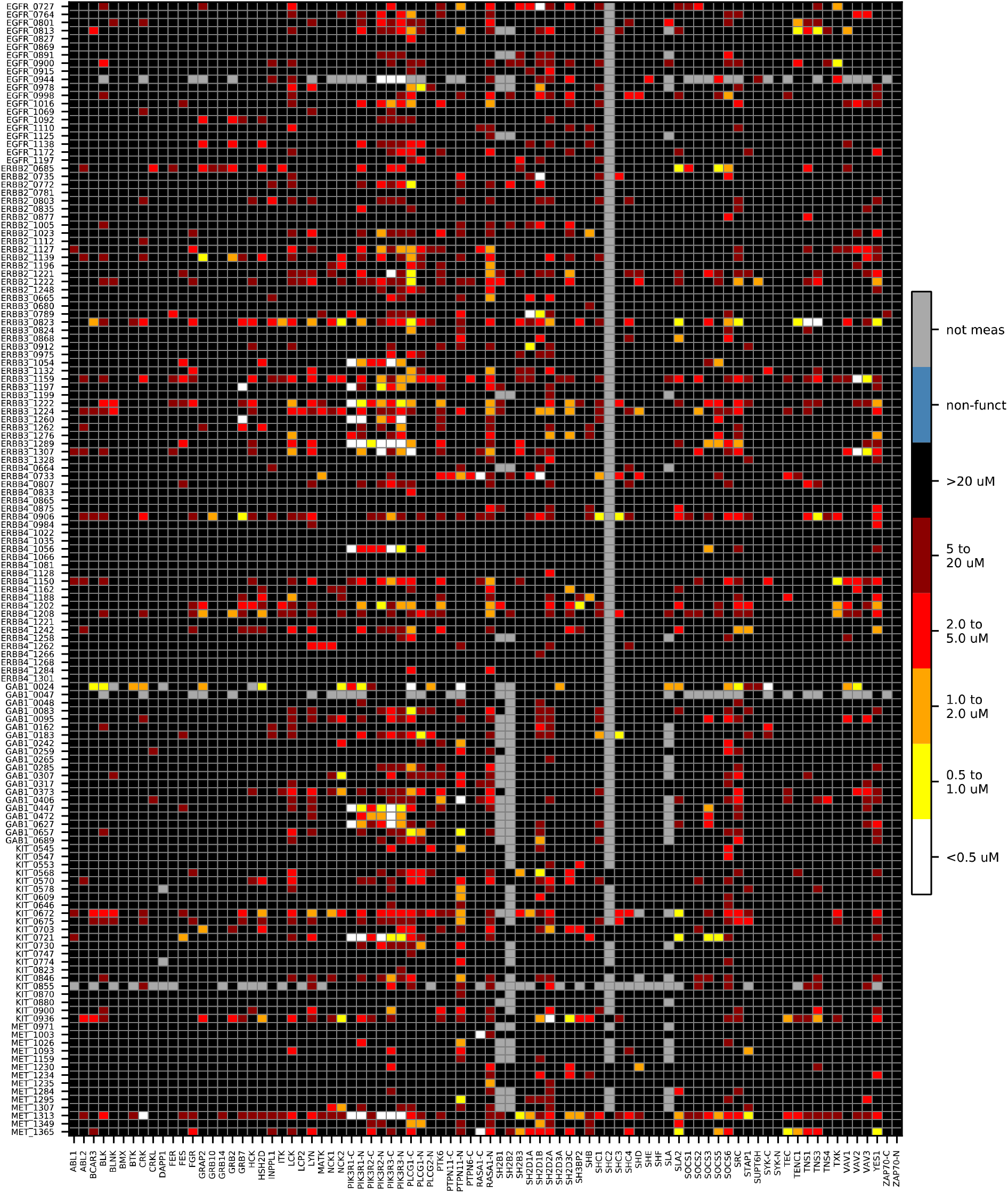
Results from the Original Publication. A heat map showing the original published results in the same format, sorting order, and naming convention – for comparison with our revised analysis.

